# Surface GluA1 and glutamatergic transmission are increased in cortical neurons of a VPS35 D620N knock-in mouse model of parkinsonism and altered by LRRK2 kinase inhibition

**DOI:** 10.1101/2021.01.18.427223

**Authors:** Chelsie A Kadgien, Anusha Kamesh, Jaskaran Khinda, Li Ping Cao, Jesse Fox, Matthew J Farrer, Austen J Milnerwood

**Affiliations:** University of British Columbia, Canada; Montreal Neurological Institute, McGill University, Canada; University of Florida, Canada

## Abstract

Vacuolar protein sorting 35 (VPS35) regulates receptor recycling from endosomes. A missense mutation in VPS35 (D620N) leads to autosomal-dominant, late-onset Parkinson’s disease. Here, we use a VPS35 D620N knock-in mouse to study the neurobiology of this mutation. In brain tissue, we confirm previous findings that the mutation results in reduced binding of VPS35 with WASH-complex member FAM21, and robustly elevated phosphorylation of the LRRK2 kinase substrate Rab10. In cultured cortical neurons, the mutation results in increased endosomal recycling protein density (VPS35-FAM21 co-clusters and Rab11 clusters), glutamate release, and GluA1 surface expression. LRRK2 kinase inhibition exerted genotype-specific effects on GluA1 surface expression, but did not impact glutamate release phenotypes. These results improve our understanding of the early effects of the D620N mutation on cellular functions that are specific to neurons. These observations provide candidate pathophysiological pathways that may drive eventual transition to late-stage parkinsonism in VPS35 families, and support a synaptopathy model of neurodegeneration.

## Introduction

VPS35 is a core component of the retromer complex, which recycles transmembrane cargoes from endosomes to the *trans*-Golgi network or surface membranes (reviewed in Bonifacino & Rojas, 2006; Burd & Cullen, 2014; Cullen & Korswagen, 2011; Seaman, Gautreau, & Billadeau, 2013). A missense mutation in VPS35 (D620N) causes late-onset autosomal Parkinson’s disease (PD) clinically indistinguishable from idiopathic PD (Struhal et al., 2014; Vilariño-Güell et al., 2011; Zimprich et al., 2011). While PD is classically thought of as a hypodopaminergic motor disorder caused by degeneration of the *substantia nigra*, recent attention has been brought to its many non-dopaminergic symptoms that often precede motor onset by years (reviewed in Beitz, 2014; Goldman & Postuma, 2014). Impaired glutamatergic plasticity in the cortex and cortical cell loss have both been observed in the brains of people with pre-motor PD (MacDonald & Halliday, 2002; reviewed in Foffani & Obeso, 2018) calling attention to glutamatergic systems. While VPS35 D620N knock-in mice (VKI) eventually develop nigral pathology (X. Chen et al., 2019), here we study fundamental alterations in VPS35 function to identify early pathways which can be targeted to potentially prevent transition to disease states.

In primary neurons, retromer clusters are dynamically active in soma, axons, dendrites, and dendritic spines (Choy et al., 2014; Munsie et al., 2015). We, and others, demonstrated that VPS35 participates in surface delivery of GluA1-containing AMPA-type glutamate receptors, supporting synapse development, maturation, and activity-dependent AMPA receptor delivery for long-term potentiation (LTP) in more mature neurons (Choy et al., 2014; Munsie et al., 2015; Tang et al., 2020; Temkin et al., 2017; Tian et al., 2015; Tsika et al., 2014; C.-L. Wang et al., 2012; Zhang et al., 2012). Exogenous expression of D620N mutant VPS35 impairs its motility in dendrites and trafficking into spines (Munsie et al., 2015). Furthermore, we showed mutant expression alters excitatory synaptic current amplitudes in mouse neurons, and AMPA receptor cluster intensities in both mouse neurons and dopamine neuron-like cells derived from human mutation carrier iPSCs (Munsie et al., 2015). Together this argues retromer is important in synapse development, maintenance, and functional connectivity. However, it is unclear how overexpression artefacts impinge on normal retromer function, and how mutations in VPS35 alter its role at the synapse.

VPS35 interacts with LRRK2, another PD-implicated protein, both physically and functionally (Inoshita et al., 2017; Linhart et al., 2014; MacLeod et al., 2013; Mir et al., 2018; Vilariño-Güell et al., 2013; Zhao et al., 2018). LRRK2 is a large multi-domain protein implicated in ~5% of all familial Parkinson’s disease through autosomal-dominant mutations and genetic risk (Healy et al., 2008). The most common LRRK2 mutation is the G2019S substitution. In neurons, LRRK2 participates in synaptic vesicle (SV) recycling and release (Beccano-Kelly, Kuhlmann, et al., 2014; Beccano-Kelly, Volta, et al., 2014; Cirnaru et al., 2014; Piccoli et al., 2011; Volta, Cataldi, et al., 2015), and in post-synaptic receptor trafficking and spine development (Matikainen-Ankney et al., 2018, 2016; Parisiadou et al., 2014; Sweet, Saunier-Rebori, Yue, & Blitzer, 2015). Inoshita and colleagues (2017) recently reported that LRRK2 localizes upstream of VPS35 at *Drosophila* neuromuscular junctions where they together regulate the SV cycle.

PD-associated mutations in LRRK2 increase its kinase activity (reviewed in Taylor & Alessi, 2020), which has been shown to phosphorylate multiple Rab-GTPases (Jeong et al., 2018; Steger et al., 2016). Increased autophosphorylation of LRRK2 and phosphorylation of Rab10 have been observed in *post-mortem substantia nigra* from individuals with idiopathic PD (Di Maio et al., 2018), monocytes from humans with the D620N mutation, and tissues from VPS35 D620N knock-in mice (Mir et al., 2018). This provides evidence that VPS35 and LRRK2 mutations converge on LRRK2 kinase, and that aberrant phosphorylation of Rab proteins involved in synaptic transmission may be related to synaptic phenotypes observed in PD models (reviewed in Kuhlmann & Milnerwood, 2020).

Here we probe early neurobiological function, potentially pathophysiological dysfunction, and drug responses in neurons from VKI mice. This may uncover factors that combine with age / environment to eventually trigger transition to pathology. We examined synapse development, structure, and function, in addition to protein levels, phosphorylation state and binding relationships in brain tissue and cultured cortical neurons from VKI mice. The D620N mutation reduced retromer complex association with its regulatory proteins, increased dendritic clustering of proteins involved in surface protein recycling, and augmented glutamate transmission. We assayed Rab10 phosphorylation in cultured cortical neurons and the effects of LRRK2 kinase inhibition on glutamatergic transmission phenotypes. LRRK2 inhibition had genotype-dependent effects on postsynaptic AMPAR trafficking and little effect on presynaptic release. In WT cells, LRRK2 kinase inhibition produced effects consistent with LRRK2 acting to regulate surface trafficking of AMPARs to developing/silent synapses.

The VPS35 knock-in mouse provides a model system in which to develop insights into the molecular and cellular effects of VPS35 mutations, and potentially the etiology PD. Furthermore, LRRK2 kinase inhibition was shown to affect post-synaptic AMPAR trafficking, and had previously unreported effects on WT synapse development that have implications for the potential utility of LRRK2 kinase inhibitors in the treatment of non-LRRK2 PD.

## Results

### Altered protein-protein binding relationships in VKI brain

Initial characterization of VKI mice revealed no alterations to VPS35, VPS26, or VPS29 protein expression in brain tissue at 3 months (Cataldi et al., 2018; X. Chen et al., 2019). In agreement, we found no genotype effect on levels of VPS35 (1-way ANOVA *p*=0.97), VPS26 Kruskal-Wallis *p*=0.73), or WASH complex member FAM21 (1-way ANOVA *p*=0.88), relative to β-tubulin in whole brain lysate from 3 month old VKI mice quantified by WES capillary-based western blot (Fig. 1 A.i-iv). Others have shown increased LRRK2-mediated phosphorylation of Rab10 at threonine 73 (pRab10) in VKI mouse brain in the absence of changes to Rab10 levels (Mir et al., 2018), a finding we replicated here using fluorescence western blot with knock-out validated phospho-specific antibodies (Fig. 1 C.i-iii; Rab10 Kruskal-Wallis *p*=0.16; pRab10 Kruskal-Wallis *p*<0.0001; Uncorrected Dunn’s Het \*\**p*<0.003, Ho \*\**p*<0.003).

**Figure 1.**
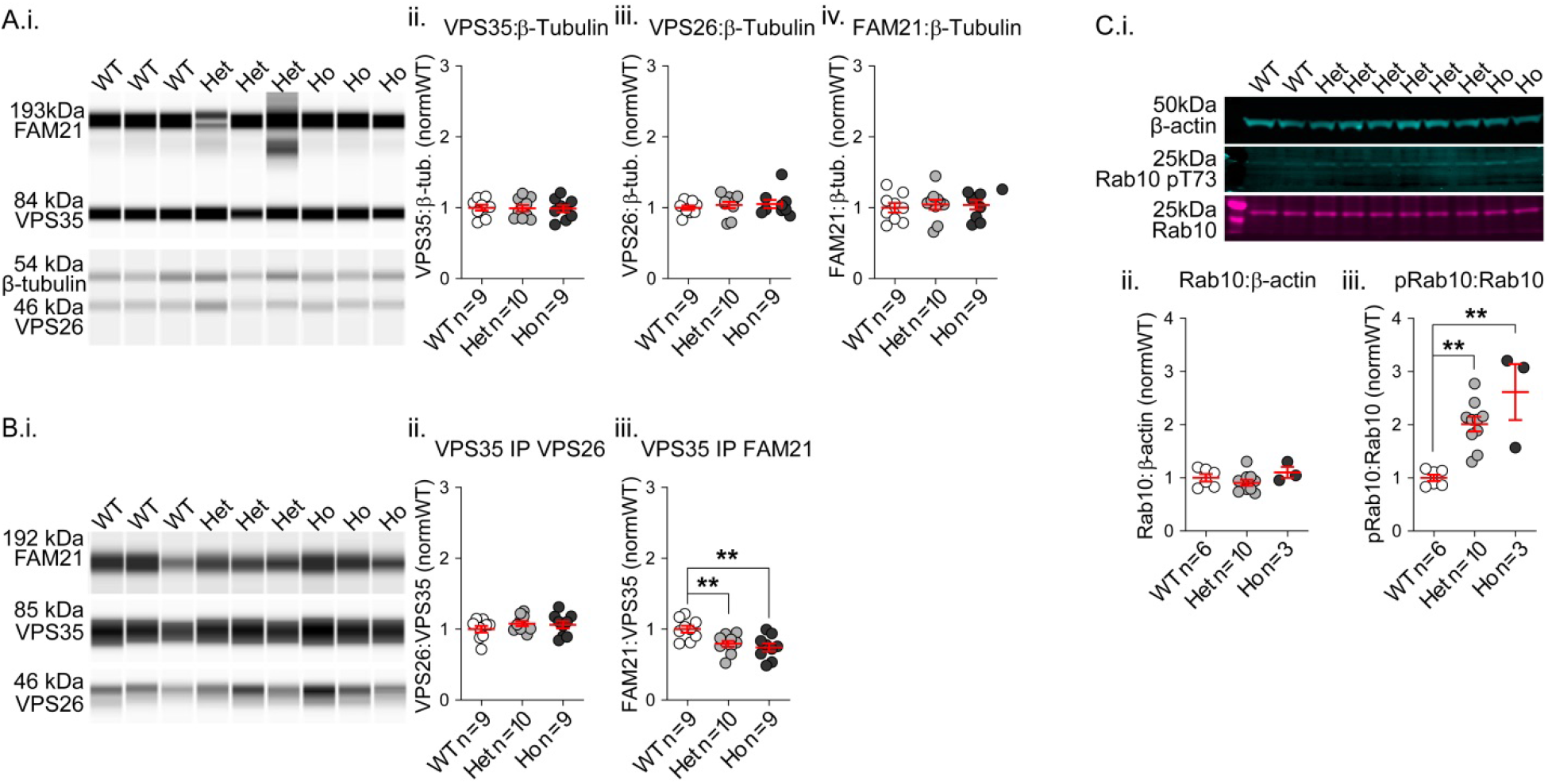
Western blot and co-immunoprecipitation of retromer and associated proteins reveals altered FAM21 binding and Rab-GTPase phosphorylation in VKI mouse brain. **A)** WES capillary-based western blot of VPS35, VPS26, FAM21, and β-tubulin in VKI whole-brain lysate (i) revealed no significant genotype effects on protein levels (ii-iv). **B)** Co-immunoprecipitations of VPS26 and FAM21 with VPS35 from VKI whole-brain lysate run on the WES system as in A (i) uncovered no effect on VPS26 pulled by VPS35 (ii) but a significant reduction in FAM21 pulled by mutant VPS35 (iii, \*\**p*<0.01). **C)** Western blot of Rab10, Rab10 phospho-T73, and β-actin in VKI whole brain lysate (i) revealed no significant genotype effect on Rab10 levels (ii), but a significant increase in Rab10 phosphorylated at threonine 73 in VKI (iii, \*\**p*<0.01).

The D620N mutation does not impair retromer trimer assembly by semi-quantitative co-immunoprecipitation (coIP) in overexpression systems (Follett et al., 2013; McGough et al., 2014; Munsie et al., 2015; Zavodszky et al., 2014) or brain lysate from VKI mice (X. Chen et al., 2019). Here we also assayed subunit assembly by coIP in brain lysate from 3-month-old VKI mice and found no genotype effect on the amount of VPS26 pulled by VPS35 (Fig. 1 Bi-ii; 1-way ANOVA *p*=0.44). In cell lines, the D620N mutation impairs VPS35-FAM21 binding (McGough et al., 2014; Zavodszky et al., 2014). In agreement, by coIP, we found the level of FAM21 pulled by VPS35 was similarly reduced in whole-brain lysates from mutant mice (Fig. 1 Bi & iii; 1-way ANOVA *p*<0.003; Uncorrected Fisher’s LSD Het \*\**p*<0.01; Ho \*\**p*<0.01).

In summary, physiological expression of VPS35 D620N does not alter protein expression levels of core retromer components, known interactor FAM21, or endolysosomal markers, but results in increased phosphorylation of the LRRK2 substrate Rab10. While retromer complex assembly appears unaltered, we found the association of VPS35 with FAM21 is reduced in VKI brain.

### No mutation effect on survival or morphology of primary cortical neurons

Neurite outgrowth in developing neurons is impaired by VPS35 knock-down (C.-L. Wang et al., 2012), and by acute overexpression of either WT or D620N VPS35 (Tsika et al., 2014). In light of this, we quantified survival and morphology of DIV21 cultured cortical neurons from VKI mice.

Genotype had no effect on cell density (Fig. 2 B; 1-way ANOVA *p*=0.47). CAG-AAV-GFP constructs were nucleofected into a subset of cortical neurons on the day of plating, and morphology was assessed at DIV21 by Sholl analysis (Fig. 2 A). There was no genotype effect on the number of intersections (Fig. 2 C; 2-way RM ANOVA radial distance x genotype *p*=0.87; genotype *p*=0.78), or secondary measures of neurite complexity (Fig. 2 D-F; Kruskal-Wallis *p*= 0.79, 0.34 & 0.29, respectively).

**Figure 2.**
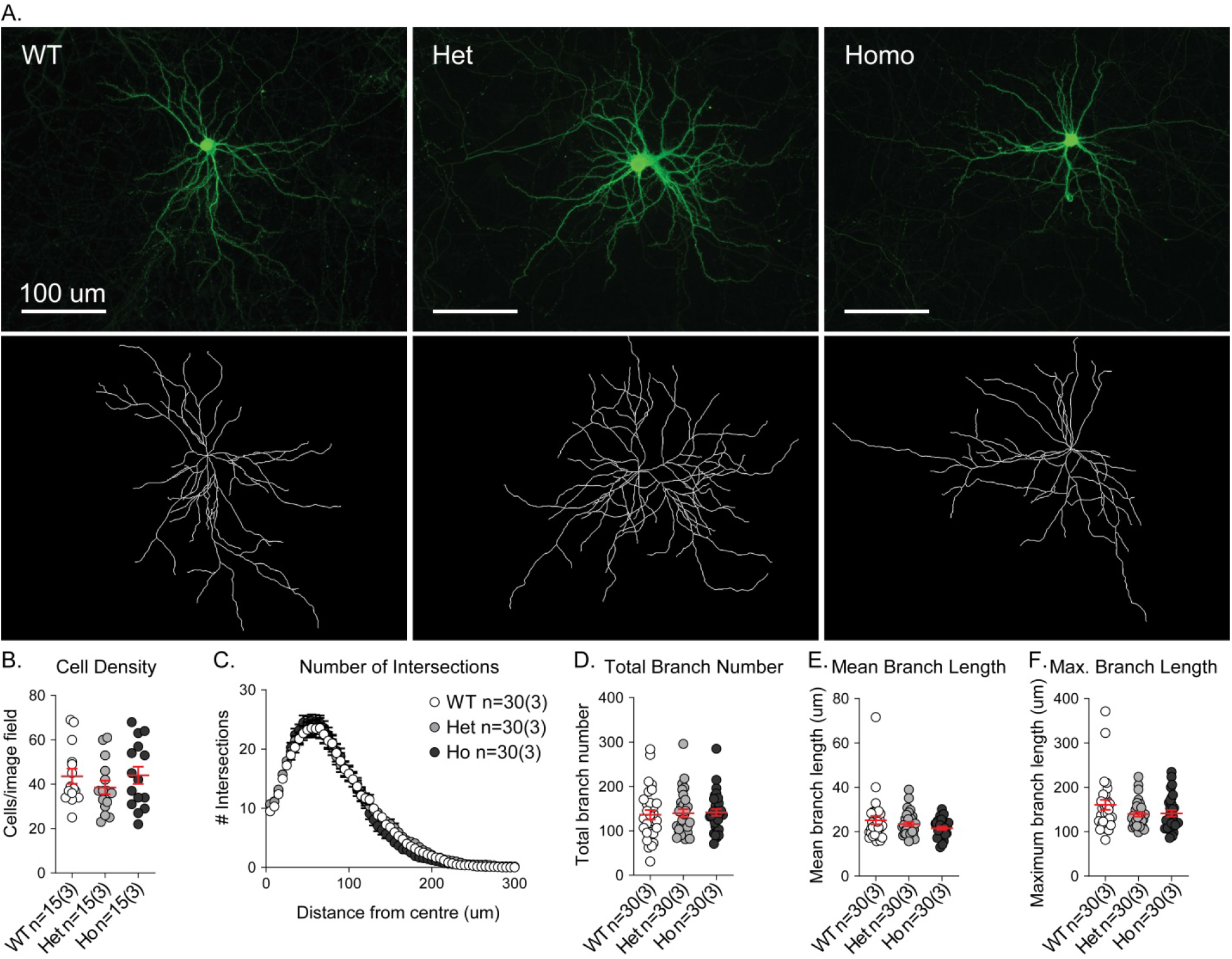
Cell density and dendritic morphology are not altered in cortical cultures from VKI mice. **A)** Cortical cells were nucleofected with CAG-AAV-GFP plasmids on the day of plating and fixed at DIV21. GFP signal was amplified and imaged (top panel), then 2D *in silico* cell reconstruction performed in ImageJ (bottom panel). **B)** There was no effect of genotype on neuron density, indicating equivalent survival and no cell death. **C)** Sholl analysis revealed no significant effect of genotype upon neurite complexity. **D-F)** There were also no genotype effects on total branch number, average branch length, or maximum branch length.

Thus, our evidence suggests that physiological expression of VPS35 D620N does not alter survival or structural development of cultured cortical neurons at 21 days *in-vitro*.

### Increased colocalization and density of endosomal recycling proteins

Since the mutation alters VPS35 interaction with FAM21, we studied the localization of VPS35, its known interactors, and endosomal markers in cortical neuron dendrites. We immunostained neurons at DIV21 to detect VPS35 with VPS26, FAM21, NEEP21, or Rab11 (Fig. 3 B-E).

**Figure 3.**
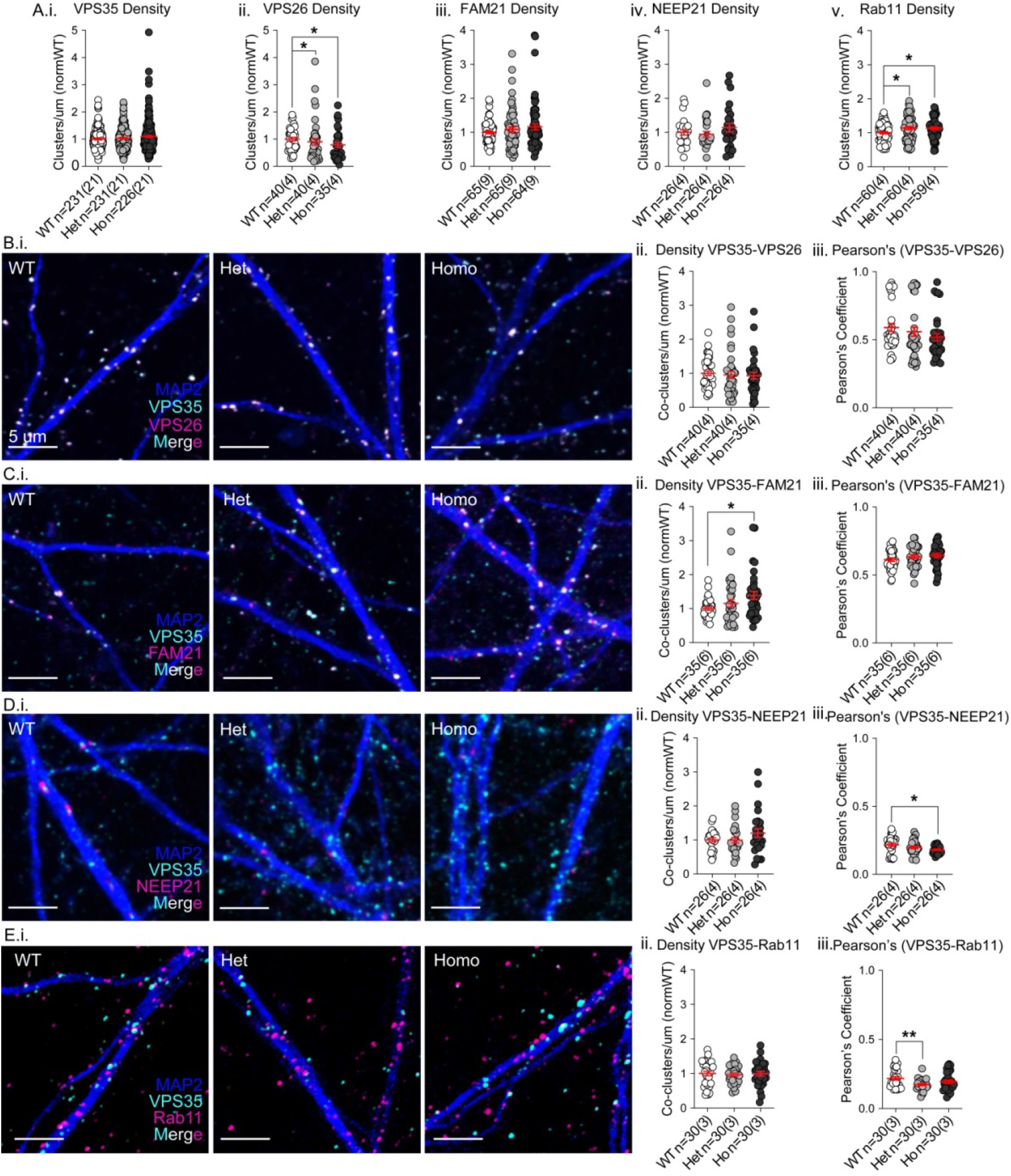
Accumulation of VPS35-FAM21 co-clusters and Rab 11 clusters in dendrites. **B-E)** Cultured cortical neurons were immunostained for MAP2 (blue), VPS35 (cyan), and VPS26, FAM21, NEEP21, or Rab11 (magenta) **A)** There was no genotype effect on dendritic cluster density of VPS35 (i), FAM21 (iii), or NEEP21 (iv). VPS26 cluster density was significantly reduced in both mutants (ii, Het \**p*<0.04 & Ho \**p*<0.02), whereas Rab11 cluster density was increased in both genotypes (v, Het \**p*<0.02 & Ho \**p*<0.03). **B)** VPS35 and VPS26 (i); there was no genotype on co-cluster density or Pearson’s coefficient (ii & iii). **C)** VPS35 and FAM21 (i); there was a significant genotype effect on co-cluster density due to increases in homozygous cells (ii, \**p*<0.02), but no genotype effect on Pearson’s coefficients (iii). **D.i)** NEEP21 and VPS35 (i); there was no genotype effect on co-cluster density (ii), but a significant genotype effect on Pearson’s coefficient due to a significant reduction in homozygous VKI dendrites (iii, \**p*<0.02). **E)** VPS35 and Rab11 (i); there was no genotype effect on the co-cluster density (ii), but a significant genotype effect in the Pearson’s coefficient due to reduced correlation in heterozygous dendrites (iii, \*\**p*<0.003).

Dendritic cluster density of VPS35 was unaffected by genotype (Fig. 3 A.i; Kruskal-Wallis *p*=0.30); however, the density of VPS26 clusters was reduced in both heterozygous and homozygous cells (Fig. 3 A.ii; Kruskal-Wallis *p*<0.04; Uncorrected Dunn’s Het \**p*<0.04 & Ho \**p*<0.02). We observed robust colocalization of VPS35 with VPS26 (Fig. 3 B.i; Pearson’s ~0.6) and no genotype effect on colocalized cluster density or Pearson’s correlation coefficient (Fig. 3 B. ii-iii; Kruskal-Wallis *p*=0.45 & 0.10, respectively).

There was no genotype effect on FAM21 cluster density (Fig. 3 A.iii; Kruskal-Wallis p=0.24). VPS35 and FAM21 were strongly colocalized (Fig. 3 C.i; Pearson’s ~0.6) in WT, with no effect of genotype on the Pearson’s coefficient (Fig. 3 C.iii; 1-way ANOVA *p*=0.24). Counterintuitive to reduced coIP with FAM21, the density of VPS35-FAM21 co-clusters was increased in mutants, in a gene dose-dependent manner, with a significant *post hoc* increase in homozygous cells (Fig. 3 C. ii; Kruskal-Wallis *p*<0.03; Uncorrected Dunn’s Ho \**p*<0.02).

Neuronal endosomal enriched protein 21 (NEEP21 or NSG1) is itinerant in dendritic endosomes and rapidly degraded with little to no recycling (Yap, Digilio, McMahon, & Winckler, 2017). We therefore used NEEP21 as a marker of the non-recycling endolysosomal pathway (Fig. 3 D.i). We observed no genotype effect on the density of NEEP21 clusters (Fig. 3 A.iv; Kruskal-Wallis *p*=0.37), nor VPS35-NEEP21 co-cluster densities (Fig. 3 D.ii; Kruskal-Wallis *p*=0.38), suggesting no change in localization to early or late endosomes. VPS35 and NEEP21 were apposed with little overlap, in accordance with retromer being active in endosomal transport for recycling, not lysosomal degradation (Fig.3 D.i; Pearson’s coefficient ~0.2); however, there was a statistically significant decrease in the Pearson’s coefficient in VKI dendrites (Fig. 3 D.iii; Welch’s ANOVA *p*<0.03; Unpaired t with Welch’s correction Ho \**p*<0.02).

Rab11 resides in recycling endosomes (REs) and participates in AMPAR surface trafficking (Bodrikov, Pauschert, Kochlamazashvili, & Stuermer, 2017; Bowen, Bourke, Hiester, Hanus, & Kennedy, 2017; Correia et al., 2008; Esteves da Silva et al., 2015; Jaafari, Henley, & Hanley, 2012; Seebohm et al., 2012), thus is was used as a marker of REs (Fig. 3 E.i). Rab11 cluster density was increased in both VKI mutants (Fig. 3 A.v; 1-way ANOVA *p*<0.03; Uncorrected Fisher’s LSD \**p*<0.02 & \**p*<0.03). Despite these increases, there was no genotype effect on the colocalization density of VPS35 and Rab11 (Fig. 3 E.ii; 1-way ANOVA *p*=0.92) but a decrease in Pearson’s coefficient in heterozygous cells (Fig. 3 E.iii; Welch’s ANOVA *p*<0.009; Unpaired t with Welch’s correction Het \*\**p*<0.003). A reduction in signal overlap in this instance may be reflective of an accumulation of recycling endosomes downstream of retromer.

VPS35 acts with the WASH complex to drive surface recycling of receptors (Derivery et al., 2009; Gomez & Billadeau, 2009), whereas Rab11 resides in recycling endosomes that collect receptors for delivery to the surface (Bodrikov et al., 2017; Bowen et al., 2017; Correia et al., 2008; Esteves da Silva et al., 2015; Jaafari et al., 2012; Seebohm et al., 2012). In summary, the D620N mutation increased clustering of proteins found on structures involved in surface recycling (Rab11) and VPS35 localization with complexes that drive forward recycling (WASH complex/FAM21). Conversely, overlap of VPS35 signal with protein found in the degradative pathway (NEEP21) was reduced. Together the observations suggest that the mutation increases capacity for surface recycling, or halts forward traffic resulting in an accumulation of Rab11 +ve recycling endosomes.

### VPS35 association with GluA1 not altered by D620N mutation

VPS35 traffics AMPA-type glutamate receptors in neurons (Choy et al., 2014; Munsie et al., 2015; Temkin et al., 2017; Tian et al., 2015; Zhang et al., 2012). We showed VPS35 interacts with GluA1 in murine brain tissue by coIP, and the mutation increased dendritic GluA1 signal intensity in ICCs of mouse primary neurons and human iPSC-derived neuron-like cells (Munsie et al., 2015).

We used western blot and coIP to investigate mutant VPS35 association with GluA1. We found no difference in GluA1 protein levels by western blot, nor coIP of GluA1 by VPS35 in 3-month-old VKI cortical lysate (Fig. 4 A.i&ii, B.i&ii; Kruskal-Wallis *p*=0.99 & 0.99, respectively). We assayed coIP of other neurotransmitter receptors in cortical and striatal lysates and discovered that VPS35 interacts with NMDA receptor subunit GluN1 and D2-type dopamine receptors (D2R), but there were no genotype effects on expression levels or co-IP (Fig. 4 Supp. 1), although this does not rule out altered traffic of these cargoes.

**Figure 4.**
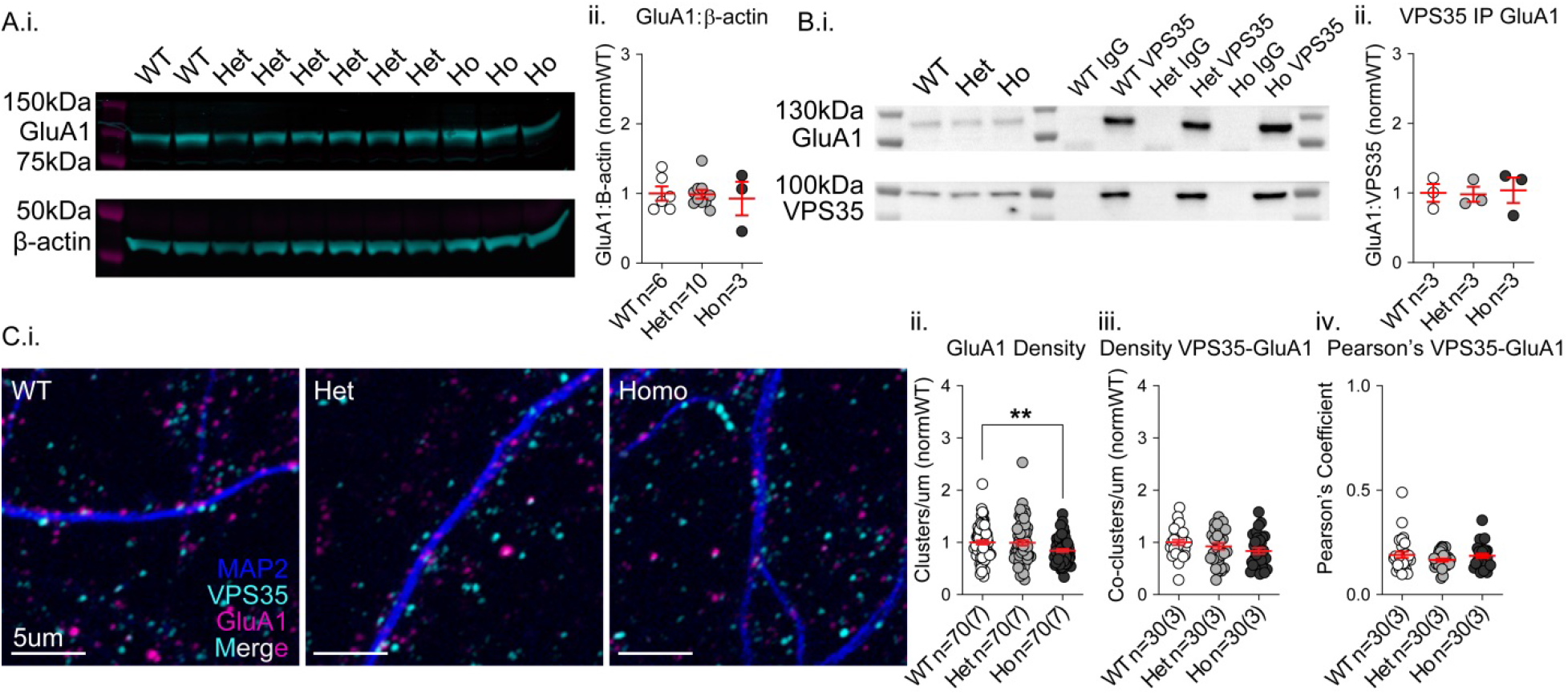
GluA1 protein expression levels are unaltered but dendritic cluster density is reduced in VKI. **A)** Western blot of GluAi and β-actin in cortical lysates of VKI mice (i) revealed no genotype effect on GluA1 protein levels (ii). **B)** Co-immunoprecipitation of GluA1 with VPS35 (i) revealed no genotype effect (ii). **C)** Cultured cortical neurons immunostained for MAP2 (blue), VPS35 (cyan), and GluA1 (magenta)(i). There was a significant reduction in GluA1 cluster density in homozygous VKI neurons (ii, \*\**p*<0.005), and no genotype effect on VPS35-GluAi co-cluster density of (iii), or Pearson’s coefficient (iv).

We performed ICC against dendritic VPS35 and GluA1 (Fig. 4 C.i) and found statistically significant reductions in dendritic GluA1 cluster density in homozygous VKI (Fig. 4 C.ii; Kruskal-Wallis *p*<0.02; Uncorrected Dunn’s Ho \*\**p*<0.005). Colocalized VPS35-GluA1 cluster density was only slightly reduced (Fig. 4 C.iii; 1-way ANOVA *p*=0.13), with little effect on the Pearson’s coefficient (Fig. 4 C.iv; Kruskal-Wallis *p*=0.57), indicating that the reduction in GluA1 clusters is primarily occurring outside of retromer-positive structures.

In summary, the D620N mutation had no effect on expression levels of GluA1 or association with retromer in cortical tissues. Dendritic GluA1 cluster density was reduced in homozygous neurons, but remaining clusters localized with VPS35 to the same extent.

### Increased glutamatergic transmission and surface AMPAR expression

VPS35 overexpression results in reduced synapse number and miniature excitatory post-synaptic current (mEPSC) frequency in cultured cortical neurons (Munsie et al., 2015). Mutant overexpression results in larger mEPSC amplitude than WT, suggesting mutant effects on AMPAR trafficking (Munsie et al., 2015). Whether this represents a gain- or loss-of-function is impossible to determine in overexpression studies. To assay the functional effects of endogenous mutant expression, we examined the number and activity of glutamatergic synapses in cortical cultures from VKI mice.

Excitatory synapses were examined by immunostaining for vesicular glutamate transporter 1 (VGluT1) and postsynaptic density protein 95 (PSD95; Fig. 5 A.i); there was no genotype effect on density of VGluT1-PSD95 co-clusters (Fig. 5 A.ii; 1-way ANOVA *p*=0.57), nor cluster density of either PSD95 or VGluT1 (Fig. 5 A.iii-iv; Welch’s ANOVA *p*=0.29 & 0.42, respectively). Alongside normal neurite morphology (Fig. 2), this suggests no differences in the density of glutamatergic synapses in VKI cultures.

**Figure 5.**
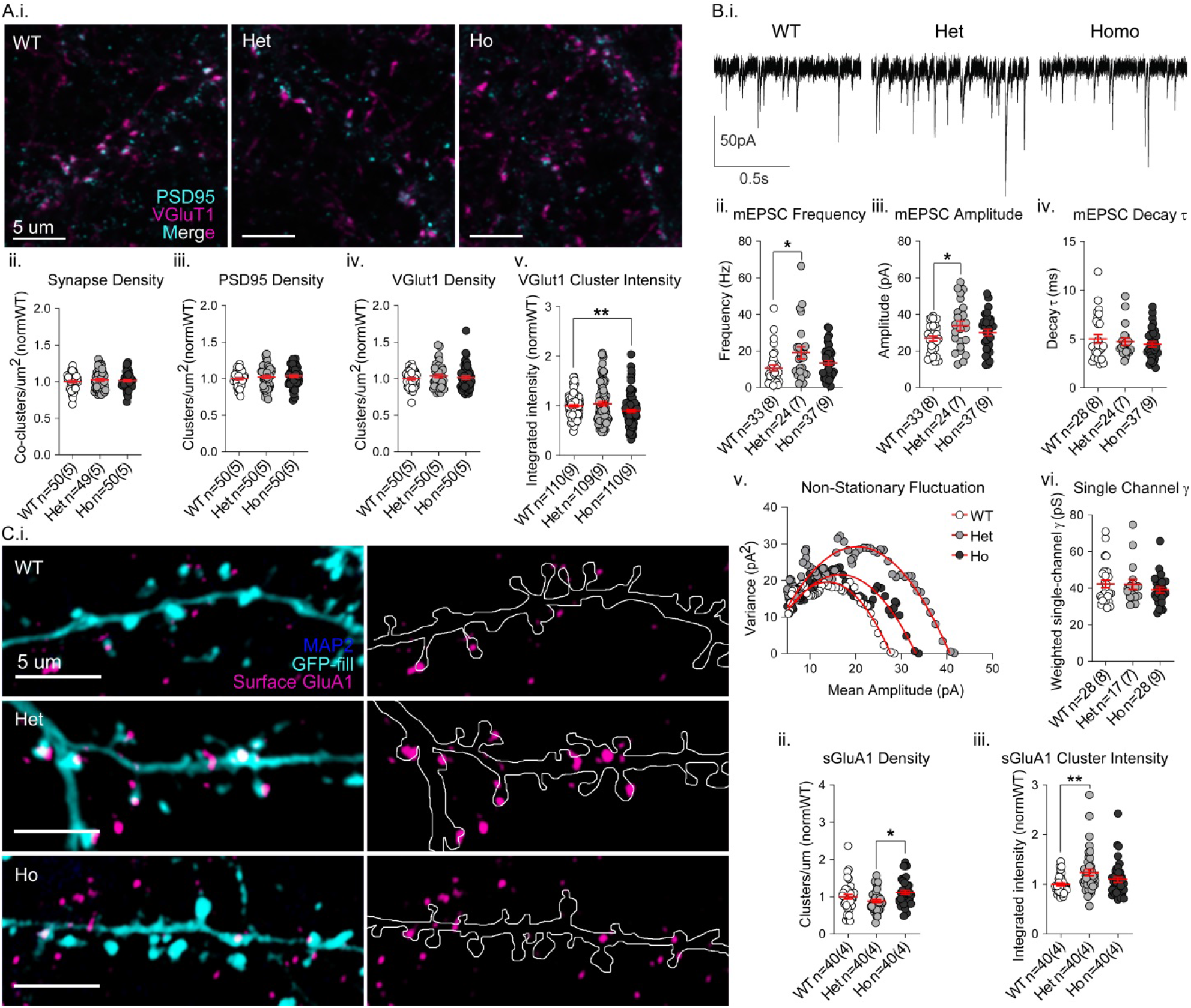
Synaptic transmission and AMPAR surface staining are increased in VKI. **A)** Cultured cortical neurons immunostained for PSD95 (cyan) and VGluT1 (magenta) (i). There was no genotype effect on synapse density defined as VGluT1-PSD95 co-clusters (ii), nor on PSD95 or VGluT1 density (iii-iv).VGluT1 cluster intensity was significantly reduced in homozygous cells (v, \*\**p*<0.003). **B)** Representative traces from whole-cell patch voltage clamp recording of mEPSCs in cortical neurons (i). There were significant genotype effects on mean mEPSC frequency (Hz) due to increases in heterozygous cells (ii, \**p*<0.02). Significant genotype effects were also seen in mEPSC amplitudes due to increases in heterozygous cells (iii, \**p*<0.02). Mean mEPSC decay times (τ) were not affected by genotype (iv). Representative mean-variance plots of mEPSC amplitude used for peak-scaled non-stationary fluctuation analysis, allowing calculation of single channel conductance (i). There was no significant effect of genotype on single channel conductance (vi, Kruskal-Wallis *p*=0.78). **C)** GFP-filled (cyan) cultured cortical cells immunostained for MAP2 (blue; to ensure no permeabilization) and surface GluA1 (magenta)(i, left panel); *in silico* neurite outlines with only GluA1 staining displayed (i, right panel). There was a significant genotype effect on surface GluA1 cluster intensity (synaptic GluA1) due to significant increases in heterozygous cells (ii, \*\**p*<0.02). There was a significant genotype effect on GluA1 cluster density, due to opposing effects on heterozygous and homozygous cells (iii, \**p*<0.05).

To assess glutamate synapse function, we performed whole-cell patch clamp recording of mEPSCs in VKI cultures (Fig. 5 B.i). Event frequency was increased, especially in heterozygous neurons (Fig. 5 B.ii; Kruskal-Wallis *p*<0.04; Uncorrected Dunn’s Het \**p*<0.02). Increased mEPSC frequency with no change in synapse density reflects either increased probability of vesicular release (Pr), or fewer AMPA-silent synapses. mEPSC amplitude was higher in mutant neurons, again most clearly in cultures from heterozygous mice (Fig. 5 B.iii; 1-way ANOVA *p*<0.05; Uncorrected Fisher’s LSD Het \**p*<0.02), suggesting increased postsynaptic AMPAR surface expression. In homozygous neurons, VGluT1 cluster intensity was decreased (Fig. 5 A.v; Kruskal-Wallis *p*<0.007; Uncorrected Dunn’s \*\**p*<0.003), possibly revealing an additional presynaptic effect in the double mutant.

Changes in amplitude may also reflect altered receptor subtype expression e.g., GluA2-lacking AMPARs which have lower channel conductance and slower kinetics (reviewed in T. Benke & Traynelis, 2019; P. Park et al., 2018), are transiently inserted during synapse unsilencing, then replaced by non-calcium permeable receptors as they mature. Thus, we examined mEPSC decay tau and single-channel conductance by peak-scaled non-stationary fluctuation analysis (NSFA), but found no significant genotype effect on either, suggesting similar AMPAR composition (Fig. 5 B.iv-vi; Kruskal-Wallis *p*=0.74 & 0.78, respectively).

To confirm increased AMPAR surface expression, we performed a surface stain against an extracellular epitope of AMPAR subunit GluA1 in non-permeabilized cells (Fig. 5 C.i). Surface GluA1 cluster intensity was higher in both mutants, significantly in heterozygous, and trending in homozygous neurons, mirroring the increases in mEPSC amplitude (Fig. 5 C.ii; Kruskal-Wallis *p*<0.007; Uncorrected Dunn’s Het \*\**p*<0.02). Neither VKI culture had surface GluA1 cluster densities significantly different from WT; however, heterozygous neurons had significantly fewer clusters than homozygous (Fig. 5 C.iii; Kruskal-Wallis *p*<0.009). Thus mEPSC frequency changes, at least in heterozygous neurons, likely reflect an increase in Pr at glutamate synapses.

Together the data demonstrate that endogenous VPS35 D620N mutations produce a gain-of-function in pre- and post-synaptic glutamate transmission. It increases postsynaptic surface AMPAR expression and mEPSC event amplitude. Frequency of mEPSC events is also increased, likely through increased presynaptic Pr. Homozygous neurons had an additional phenotype of reduced VGluT1 intensity and dendritic GluA1 cluster density.

### LRRK2 kinase inhibition affects postsynaptic AMPAR trafficking in cortical neurons

LRRK2-mediated phosphorylation of Rab10 (pRab10) is increased in brain tissue from VPS35 D620N knock-in mice (Fig. 1), in agreement with Mir *et al.* (2018). Here we also found that acute treatment with the LRRK2 kinase inhibitor MLi-2 reversed pRab10 increases in whole brain lysate from VKI mice without affecting expression levels of LRRK2, Rab10, VPS35, GluA1, or VGluT1 (Fig. 6 Supp. 1).

Synaptic phenotypes in LRRK2 G2019S knock-in mice are responsive to LRRK2 kinase inhibition (Matikainen-Ankney et al., 2016), thus we hypothesized that MLi-2 would rescue glutamatergic phenotypes in primary cortical cultures from VKI animals. We treated DIV21 primary cortical cultures for 2 hours with 500nM MLi-2. LRRK2 and LRRK2 pS935 (pLRRK2) were detected in culture lysates, and pLRRK2 appeared reduced after MLi2 treatment in all genotypes, but we deemed bands too close to background to reliably quantify (Fig. 6 Supp. 2). As a proxy for LRRK2 activity *in vitro*, we found mutation dose-dependent increases in pRab10 (as seen in brain) and a significant treatment effect in VKI (Fig. 6 A.ii 2-way ANOVA genotype x treatment *p*<0.08; genotype *p*<0.03; treatment *p*<0.0004; Uncorrected Fisher’s LSD WT-WTML12 *p*=0.99; Het-HetML12 *p*=0.26; Ho-HoML12 \*\**p*<0.005). Thus, D620N results in increased pRab10 in cultured cortical neurons at DIV21, which is reversed by treatment with MLi-2.

**Figure 6.**
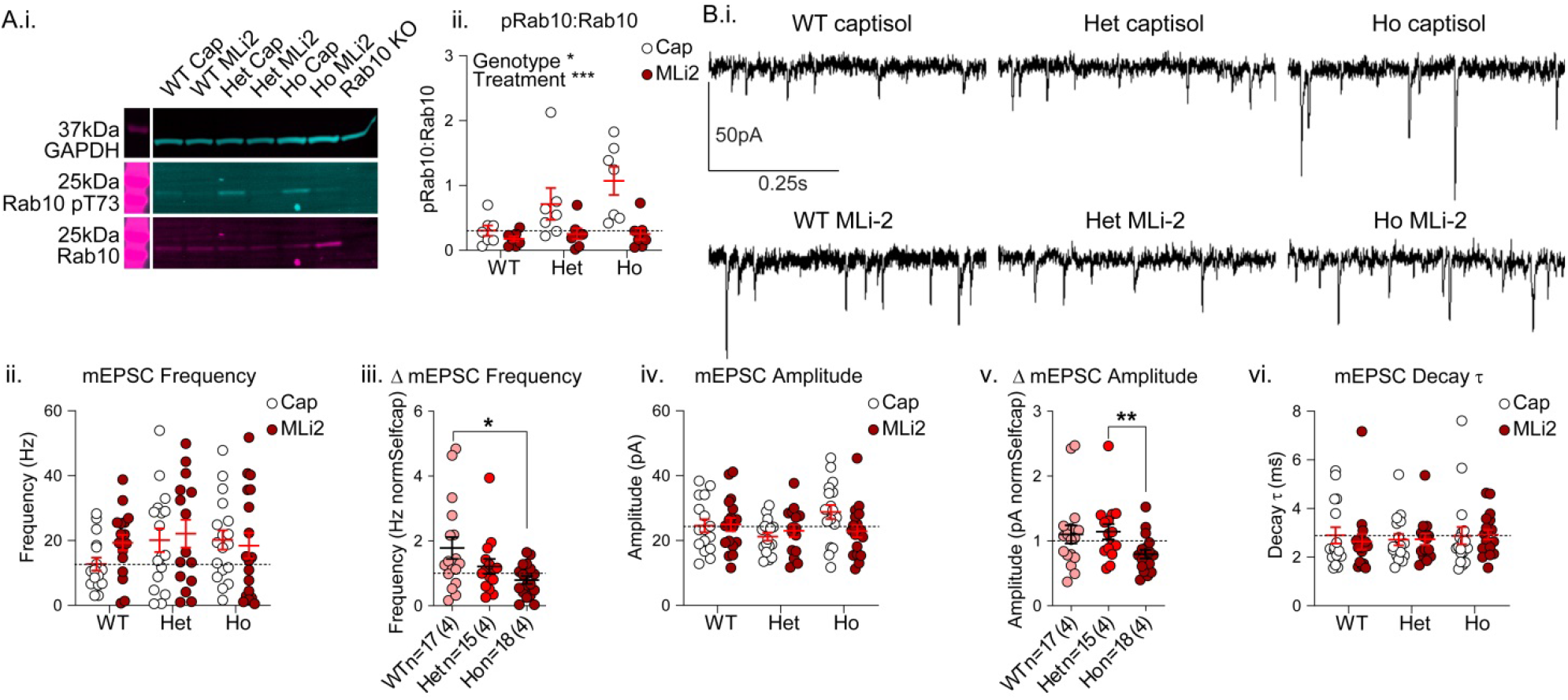
MLi-2 treatment inhibits LRRK2 kinase activity and has genotype-dependent effects on glutamate transmission. **A)** Fluorescence western blots of MLi-2 or vehicle treated cortical culture lysates were probed for Rab10, Rab10 phospho-T73 (pRab10), and GAPDH (i). There were significant genotype and treatment effects on pRab10 (ii, a proxy for LRRK2 kinase activity) due to significant increases in pRab10 in vehicle-treated homozygous neurons over WT (\**p*<0.01), which are significantly reduced by treatment (\*\**p*<0.005). **B)** Representative traces from whole-cell patch voltage clamp recording of mEPSCs in cortical neurons following acute MLi-2 treatment (i). There were no significant genotype or treatment effects on mean mEPSC frequency (Hz) (ii); however, normalization of the treatment effect in each genotype to its own control revealed significant genotypes on treatment effects, due to a nearly two-fold increase in mEPSC frequency in only WT cells following treatment (iii, \*\**p*= 0.01). There were no statistically significant genotype or treatment effects on mEPSC amplitude (iv), but similar within-genotype normalization revealed significantly different treatment effects in heterozygous and homozygous neurons were statistically different due mildly opposing effects in heterozygous and homozygous cultures (v, \*\**p*<0.01). There were no genotype or treatment effects on mEPSC decay times (τ). For Aii, n=7 independent cultures for all groups. For Bii, iv, vi: WTCap n=16(4), WTMLi2 n=17(4), HetCap n=17(4), HetMLi2 n=15(4), HoCap n=18(4), HoMLi2 n=18(4).

We next investigated the effect of acute MLi-2 treatment on synapse function by whole-cell patch clamp. We found no statistically significant differences in mEPSC frequency (Fig. 6 B.ii); although event frequency was generally higher in vehicle treated mutants relative to WT (in agreement with basal increases; Fig.5), this higher frequency was not affected by MLi-2 treatment (2-way ANOVA interaction *p*=0.44; genotype *p*=31; treatment *p*=0.40). We observed an unexpected 2-fold increase in mEPSC frequency in treated WT cells. Thus, the genotype-normalized drug effects show that MLi-2 treatment increased mEPSC frequency in WT cells and had no effect in heterozygous or homozygous neurons (Fig. 6 B.iii; Kruskal-Wallis *p*<0.04; Uncorrected Dunn’s WT-Ho \*\**p*< 0.01).

The basal increases in mEPSC amplitude in heterozygous mutants (Fig 5) were seemingly lost in vehicle. Amplitude was higher in homozygous controls compared to heterozygous; however the two-way ANOVA failed to reach statistical significance (Fig. 6 B.iv; 2-way ANOVA interaction *p*=0.10; genotype *p*=0.14; treatment *p*=0.41). MLi-2 increased amplitude slightly in WT and heterozygous cells and produced a significantly different effect (a reduction) in homozygous cells (Fig. 6 B.v; Kruskal-Wallis *p*<0.03; Uncorrected Dunn’s Het-Ho \*\**p*<0.01). We observed no differences in mEPSC decay tau of vehicle or MLi-2 treated cultures (Fig. 6 B.vi; 2-way ANOVA genotype x treatment *p*=0.83; genotype *p*=0.81; treatment *p*=0.77), suggesting no alteration to AMPAR subunit composition.

To further investigate effects of LRRK2 inhibition in WT & VKI cultures, we performed immunostaining for synapses and surface GluA1. There were no statistically significant effects of genotype or treatment on excitatory synapse density (Fig. 7 A.iv; 2-way ANOVA genotype x treatment *p*=0.08; genotype *p*=0.52; treatment *p=*0.68), despite the trending increase in vehicle-treated mutants, compared to WT. There was a significant genotype x treatment interaction effect on both PSD95 (Fig. 7 A.ii; 2-way ANOVA genotype x treatment *p*<0.03; genotype *p*=0.72; treatment *p=*0.22; Uncorrected Fisher’s LSD WT-Het \**p*<0.05; Het-HetMLi2 \**p*<0.04) and VGluT1 (Fig. 7 A.iii; 2-way ANOVA genotype x treatment *p*<0.03; genotype *p*=0.47; treatment *p*=0.67; Uncorrected Fisher’s LSD Ho-HoMLi2 \**p*<0.03) cluster densities, seemingly due to MLi-2 decreasing PSD95 cluster density in heterozygotes, and VGluT1 density in homozygotes. Differences in synapse density are in the same pattern as frequency changes; however, their magnitude is lower, suggesting they are unlikely the primary source of frequency differences.

**Figure 7.**
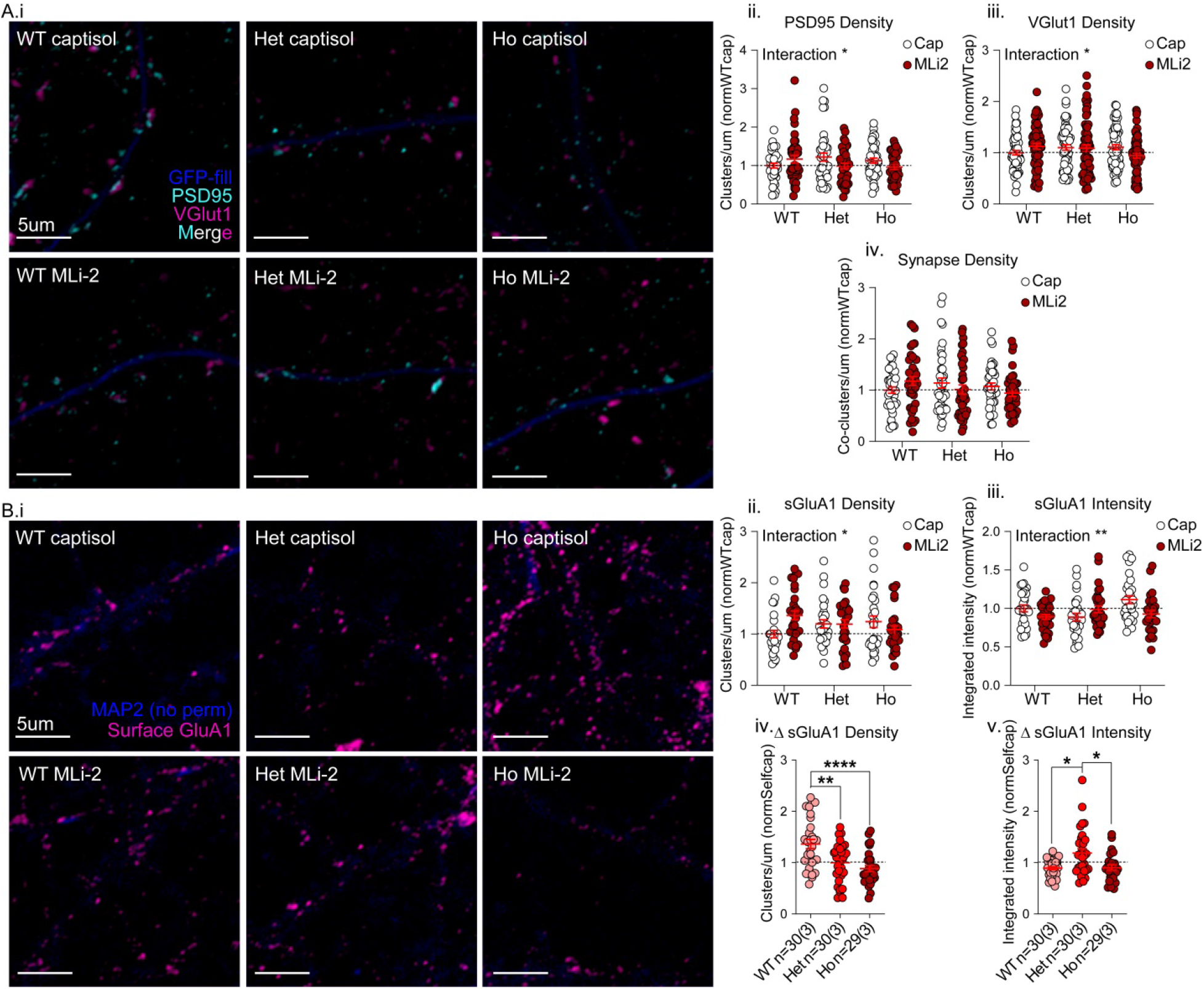
MLi-2 treatment alters localization of synaptic proteins in a genotype-dependent manner. **A)** Cultured cortical neurons immunostained for MAP2 (blue), PSD95 (cyan), and VGluT1 (magenta) following acute MLi-2 or vehicle treatment (i). There were interaction effects on PSD95 density due to increases in PSD95 density in heterozygous controls over WT (ii, *p<0.05) and a reduction in heterozygous neurons treated with MLi-2 *p<0.04. There were also interaction effects on VGluT1 density due to a reduction in cluster density in homozygous cells following MLi-2 treatment (iii, \**p*=0.03); however, there were no statistically significant effects on synapse (PSD95-VGluT1 co-cluster) density (iv). **B)** Non-permeabilized cultured cortical cells immunostained for MAP2 (blue), and surface GluA1 (magenta) following acute MLi-2 or vehicle treatment (i). There was a significant interaction effect on surface GluA1 cluster density (ii) due to an increase in WT surface GlulA1 clusters (\*\**p*<0.004); There was a significant interaction effect on surface GluA1 intensity (iii), though in this case it was due to underlying differences in control heterozygous and homozygous cells (\*\*\**p*<0.0004) and a treatment effect in homozygous cells (\*\**p*<0.004). Normalization of treatment effects within genotype revealed that changes to GluA1 surface density (iv) were significantly larger in WT cells than heterozygous (\*\**p*<0.003) and homozygous mutant cells (\*\*\*\**p*<0.0001), and that treatment had opposite effects on GluA1 intensity (v) in heterozygous cells than it did in WT (\**p*<0.02) and homozygous cells (\**p*<0.02). For Aii-v, n=40(4) for all groups and for Bii-iii, n=30(3).

Surface GluA1 cluster density revealed a significant genotype x treatment interaction (Fig. 7 B.ii; 2-way ANOVA genotype x treatment *p*<0.02; genotype *p*=0.93; treatment *p*=0.34; Uncorrected Fisher’s LSD WT-WTMLi2 \*\**p*<0.004), due to MLi-2 producing a, ~50% increase in WT cluster density, but having no effect in mutants (Fig. 7 B.iv; Welch’s ANOVA *p*<0.0002; Unpaired t with Welch’s correction WT-Het \*\**p*<0.003; WT-Ho \*\*\*\**p*<0.0001). The effect of MLi-2 on surface GluA1 cluster density and mEPSC frequency (with minor changes in synapse density), in WT cells suggests LRRK2 kinase inhibition may result in delivery of AMPARs to a small number of new synapses and more so to existing AMPA-silent synapses.

In vehicle treated cells, surface GluA1 cluster intensity appeared to no longer be increased in heterozygotes (compared to untreated cells in Fig.5). Generally, surface GluA1 cluster intensity closely resembles the mEPSC amplitude data and there was a statistically significant interaction effect, stemming from increased intensity in vehicle-treated homozygous cells (relative to heterozygous) and MLi-2 inducing a reduction in homozygous mutants (Fig. 7 B.iii; 2-way ANOVA genotype x treatment *p*<0.005; genotype *p*=0.13; treatment *p*=0.08; Uncorrected Fisher’s LSD Het-Ho \*\*\**p*<0.0004; Ho-HoMLi2 \*\**p*<0.004). Thus, similar to mEPSC amplitude, MLi-2 treatment resulted in different effects on surface GluA1 cluster intensity in heterozygous cells (increase), than in WT and homozygous cells (decrease; Fig. 7 B.v; Kruskal-Wallis *p*<0.02; WT-Het \**p*<0.02; Het-Ho \**p*<0.02), suggesting a genotype-dependent effect of LRRK2 kinase inhibition on AMPAR surface expression, or an interaction with the apparent effect of the vehicle Captisol.

Rab10 is involved in actin dynamics at recycling endosomes (P. Wang et al., 2016) and transport of retromer cargoes including GLR-1 (Alshafie et al., 2020; Bruno, Brumfield, Chaudhary, Iaea, & McGraw, 2016; Y. Chen et al., 2005; Glodowski, Chen, Schaefer, Grant, & Rongo, 2007; Pan, Zaarur, Singh, Morin, & Kandror, 2017). Given that pRab10 was increased in mutant cultures (Fig. 6) and that MLi-2 affected surface AMPARs (Fig.7), we analyzed dendritic Rab10 colocalization with VPS35 and GluA1 in cultured cortical neurons from VKI mice at DIV21 by ICC. We found no significant genotype effect on Rab10 cluster intensity (Fig. 8 A.i; Kruskal-Wallis *p*=0.66), but a strong trend to increased Rab10 cluster density in both mutants (Fig. 8 A.ii; Kruskal-Wallis *p*<0.06). However, co-cluster densities of VPS35-Rab10 and GluA1-Rab10 were low, along with correspondingly low Pearson’s coefficients (Fig. 8 B.i&C.i; Pearson’s ~0.18 & 0.17, respectively). Although there was a trend toward increased co-cluster density in heterozygous cells in both cases (Fig. 8 B.ii&C.ii; 1-way ANOVA *p*=0.18 & 0.24, respectively) there were no genotype effects on Pearson’s coefficients for VPS35-Rab10, nor GluA1-Rab10 (Fig. 8 B.iii&C.iii; Kruskal-Wallis *p*=0.38 & 0.54, respectively). The data demonstrate that Rab10 clusters may be increased along VKI dendrites, but do not provide any evidence that Rab10 is involved in local surface delivery or recycling of AMPARs in dendrites.

**Figure 8.**
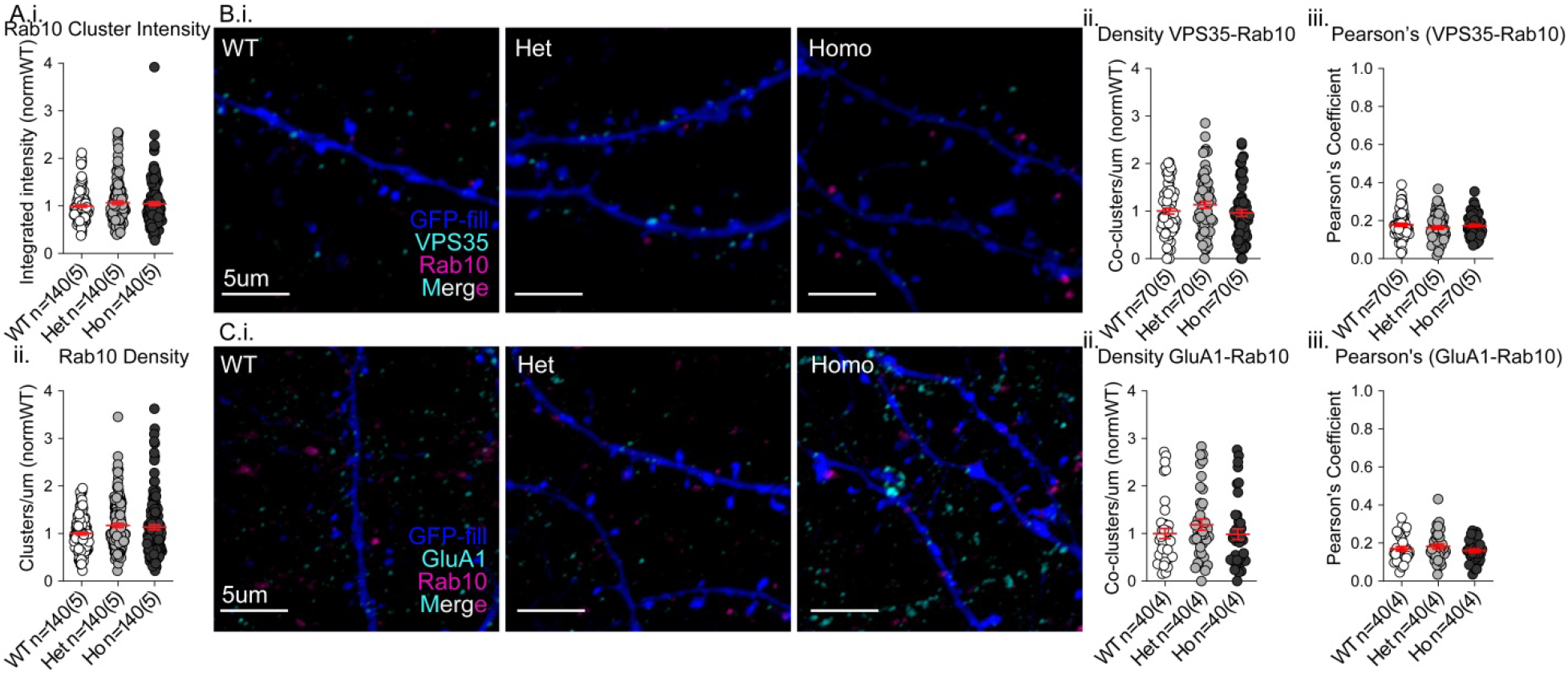
Rab10 does not colocalize with VPS35 or GluA1 in cortical neurites. **A)** There was no genotype effect on Rab10 cluster intensity (i); however, Rab10 cluster density was increased in both mutant genotypes, falling just shy of statistical significance (ii, *p*<0.06). **B)** GFP-filled (blue) cortical neurons immunostained for Rab10 (magenta), and VPS35 (cyan)(i). There were no genotype effects on co-cluster density (ii) or Pearson’s coefficient (ii). **C)** GFP-filled (blue) cortical neurons immunostained for Rab10 (magenta), and GluA1 (cyan)(i). There were no genotype effects on co-cluster density or Pearson’s coefficient (ii-iii).

Together, these experiments suggest MLi-2 treatment in WT neurons increases mEPSC frequency through delivery of AMPARs to silent synapses. MLi-2 did not rescue elevated frequency in mutants. There was no significant effect of MLi-2 on WT mEPSC amplitudes or surface GluA1 cluster intensity. MLi-2 had opposite effects in the two mutant conditions: increasing mEPSC amplitudes and surface GluA1 in heterozygous cells, and rescuing elevated mEPSC amplitudes and surface GluA1 expression in homozygous cells. Together the data suggests that LRRK2 kinase inhibition affects glutamatergic transmission in WT and VPS35 D620N mutant neurons, primarily by altering postsynaptic AMPAR trafficking, but had a clear rescue effect only on homozygous post-synaptic phenotypes (mEPSC amplitude and surface GluA1 intensity).

## Discussion

The D620N mutation did not affect expression levels of retromer components, accessory proteins, or cargoes in brain tissue from VKI mice. Retromer assembly and association with cargo by coIP was unaffected, whereas association of FAM21 was reduced by 25-30%. In cortical neuron dendrites, the mutation increased the density of clustered FAM21 colocalized with VPS35, and increased the density of Rab11 clusters. Surface expression of GluA1 was increased, alongside higher mEPSC amplitudes and frequencies with no change in excitatory synapse number, suggesting the mutation results in a gain-of-function in both AMPAR trafficking and glutamate release. We replicated findings that the D620N mutation increases LRRK2-mediated phosphorylation of Rab10. However, inhibition of LRRK2 with MLi-2 had no effect on glutamate release, while producing genotype-dependent effects on AMPAR surface expression. This suggests that mutant Pr phenotypes are LRRK2 independent, whereas LRRK2 regulates AMPAR surface traffic.

VPS35 is implicated in trafficking GluA1-containing AMPARs in overexpression and knock-down/out studies (Choy et al., 2014; Munsie et al., 2015; Temkin et al., 2017; Tian et al., 2015; Vazquez-Sanchez, Bobeldijk, Dekker, Van Keimpema, & Van Weering, 2018). Successful surface delivery of retromer cargo requires the WASH complex (including FAM21) to regulate endosomal actin dynamics, and the auxiliary protein SNX27 for correct targeting (Derivery, Helfer, Henriot, & Gautreau, 2012; Derivery et al., 2009; Gomez & Billadeau, 2009; Lee, Chang, & Blackstone, 2016; Temkin et al., 2011). Consistent with other reports (McGough et al., 2014; Zavodszky et al., 2014), we confirm a mutation-dependent reduction in VPS35 and FAM21 interaction by coIP in brain tissue from VKI mice, in the absence of changes to retromer assembly or protein levels (X. Chen et al., 2019; Follett et al., 2013; McGough et al., 2014; Munsie et al., 2015; Tsika et al., 2014; Zavodszky et al., 2014). This is in contrast to a recent report of no detectable interaction between VPS35 and FAM21 in murine brain (X. Chen et al., 2019); here we used the same extraction buffer, animal age, and tissue type, suggesting protein-protein interactions may have been disrupted in the previous study by differences in other steps of the lysis protocol. Furthermore, we observed robust colocalization of FAM21 with VPS35 in the dendrites of cultured cortical neurons.

Given the reduced association of FAM21 by coIP, the observed increase in VPS35 and FAM21 cluster colocalization and surface AMPAR expression are puzzling. The data suggests that, at least in dendritic endosomes, VPS35 D620N results in increased recruitment or retention of the WASH complex. The WASH complex is thought to associate with retromer via direct binding of the C-terminal tail of FAM21 with VPS35 (Gomez & Billadeau, 2009; Harbour, Breusegem, & Seaman, 2012; Jia, Gomez, Billadeau, & Rosen, 2012), and with SNX27 (Lee et al., 2016). The mutation may reduce association of FAM21 with VPS35 (reduced coIP), yet still allow its recruitment to surface-bound retromer in dendrites through the SNX27 interaction. Other reports have shown FAM21 acts downstream of receptor sorting, by ending WASH complex activity (Lee et al., 2016; L. Park et al., 2013). In this light, reduced (but not ablated) binding of FAM21 may prolong WASH complex-mediated receptor sorting into recycling pathways.

It is noteworthy that binding differences and accumulation of recycling proteins occurred in either a gene-dose-dependent manner or equally in the mutant neurons, whereas GluA1 surface expression and mEPSC amplitudes were more elevated in heterozygous than homozygous neurons. The observed reduction in presynaptic VGluT1 cluster intensity and postsynaptic GluA1 cluster density in only homozygous neurons suggest that compensatory mechanisms may be occurring that either do not occur in heterozygotes, or that do so at different time points.

Further to postsynaptic effects on AMPA receptors, we observed increased presynaptic glutamate release, evidenced by increased mEPSC frequency with no change in synapse number or surface GluA1 density. Other proteins associated with PD participate in the SV cycle. Alpha-synuclein, which confers the second highest genetic risk for PD, is a presynaptic regulatory protein associated with endosomal and vesicular membranes (Bendor, Logan, & Edwards, 2013; Boassa et al., 2013; Logan, Bendor, Toupin, Thorn, & Edwards, 2017; Westphal & Chandra, 2013). LRRK2 is similarly endosomally localized (Schreij et al., 2015), co-immunoprecipitates with VPS35 (MacLeod et al., 2013; Vilariño-Güell et al., 2013), and regulates presynaptic release of glutamate and dopamine (Beccano-Kelly, Kuhlmann, et al., 2014; Beccano-Kelly, Volta, et al., 2014; Cirnaru et al., 2014; Matikainen-Ankney et al., 2016; Piccoli et al., 2011; Volta, Cataldi, et al., 2015).

To date, little is known about retromer activity in the presynapse. Retromer is highly mobile in axons and present in glutamatergic synaptic boutons in cultured murine neurons (Munsie et al., 2015; Vazquez-Sanchez et al., 2018), but retromer deficiency or acute knock-down in murine hippocampal slices has no effect on synaptic glutamate release (Temkin et al., 2017; Tian et al., 2015), or SV exo- or endocytosis (Vazquez-Sanchez et al., 2018). Mutant VPS35 overexpression in cultured murine cortical cells does reduce mEPSC frequency, but also synapse number, as does WT overexpression (Munsie et al., 2015). There is no evidence thus far that retromer has a critical role in glutamate release *per se*; however, retromer associates with presynaptically expressed transmembrane proteins such as D2R (present study) and DAT (C. Wang et al., 2016). Thus retromer may be involved in transmitter release indirectly by regulating axonal transport and/or recycling of synaptic proteins that modulate release.

While much remains to be discovered about presynaptic retromer function, it is noteworthy that PD-associated mutations in LRRK2 cause similar increases in neurotransmitter release (Beccano-Kelly, Kuhlmann, et al., 2014; Matikainen-Ankney et al., 2016; Sweet et al., 2015; Volta et al., 2017; Yue et al., 2015), which are rescued by inhibition of LRRK2 kinase (Matikainen-Ankney et al., 2016; Sweet et al., 2015). Furthermore, LRRK2 mutant expression also has effects on postsynaptic processes (Matikainen-Ankney et al., 2018, 2016; Parisiadou et al., 2014; Sweet et al., 2015), indicating that, like VPS35, LRRK2 may be active in both pre- and postsynaptic compartments. Here we replicated recent reports of increased LRRK2 kinase activity in VKI mouse brain (Mir et al., 2018). Thus, we set out to test whether glutamate transmission phenotypes observed here are rescued by LRRK2 kinase inhibition.

In WT neurons, MLi-2 increased the frequency of mEPSCs nearly two-fold, likely related to the increased density of surface GluA1 clusters. GluA1-containing AMPARs are delivered to quiescent synapses during development and synaptic plasticity (reviewed in T. Benke & Traynelis, 2019; Diering & Huganir, 2018; P. Park et al., 2018; Purkey & Dell’Acqua, 2020). These data imply that inhibition of LRRK2 kinase activity results in AMPAR delivery to silent synapses in WT. In other reports, LRRK2 kinase inhibition in WT animals, or LRRK2 kinase-dead mutant expression, had no effect on neurotransmitter release (Matikainen-Ankney et al., 2016; Sweet et al., 2015), consistent with changes observed here occurring through postsynaptic mechanisms.

Similarly, we observed no effect of MLi-2 on spontaneous glutamate transmission in either mutant, as evidenced by no change in the frequency of mEPSCs or synapse density. The benefits of MLi-2 on postsynaptic receptor expression in mutant cells are enigmatic, in part due to genotype-dependent effects of the vehicle, Captisol. In heterozygous neurons, MLi-2 increased mEPSC amplitude and surface GluA1 intensity, with no change in surface GluA1 cluster density. Conversely, in homozygous cells with large mEPSC amplitudes and bright GluA1 clusters, LRRK2 kinase inhibition reduced these to below WT control levels. It is important to note here that most human D620N carriers are heterozygous for the mutation.

While the effectiveness of MLi-2 treatment for rescuing D620N phenotypes is muddied by vehicle effects, it is clear LRRK2 kinase inhibition impacted AMPAR traffic. We do not yet know if LRRK2 participates in recycling, or trafficking of newly synthesized receptors, nor the organelle loci of regulation. In cell lines, LRRK2 localizes to early endosomes (Schreij et al., 2015), phosphorylates Rab5 (Jeong et al., 2018; Steger et al., 2016), and can regulate activation of Rab7 (Gómez-Suaga et al., 2014), a Rab-GTPase involved in retromer trafficking (Rojas et al., 2008; Seaman, Harbour, Tattersall, Read, & Bright, 2009). Rab7 activation is also impaired by Rab10 knock-down, suggesting that Rab10 may act upstream in endosomal recycling (Rivero-Ríos, Romo-Lozano, Fernández, Fdez, & Hilfiker, 2020); however, we did not observe dendritic colocalization of Rab10 with either VPS35 or GluA1, suggesting that if Rab10 is involved, it is not participating locally in dendrites. Overexpression studies have also shown LRRK2 localized in the *trans-Golgi* network (Alexandria Beilina et al., 2014; Liu et al., 2018; MacLeod et al., 2013; Purlyte et al., 2018) alongside inactive Rab10 (Liu et al., 2018; Purlyte et al., 2018), thus it remains possible that LRRK2 and Rab10 regulate trafficking of newly synthesized GluA1 from the soma.

However, Rab10 is not the only substrate of LRRK2; LRRK2 may act in AMPAR trafficking through a different substrate (e.g., Rab5, Rab8, NSF), or indirectly via PKA (Parisiadou et al., 2014). While LRRK2 and VPS35 do not colocalize in immortalized cells (Alexandra Beilina et al., 2020; Alexandria Beilina et al., 2014), they do coIP together in brain lysate (MacLeod et al., 2013; Vilariño-Güell et al., 2013), suggesting they may interact in neuron-specific pathways yet to be fully described. While the site of interaction and underlying mechanism are still unclear, we provide evidence for a postsynaptic function of LRRK2 in the regulation of AMPAR surface expression, and propose that LRRK2 kinase activity acts as a tuning mechanism for synaptic strength.

In cortical cultures from LRRK2 G2019S knock-in mice, Pr is also increased, but mEPSC amplitudes are not (Beccano-Kelly, Kuhlmann, et al., 2014). Glutamate release phenotypes in *ex vivo* slices from young LRRK2 knock-in mice are rescued by MLi-2 (Matikainen-Ankney et al., 2016). Here we observe effects of VPS35 D620N on both pre- and postsynaptic processes, and an effect of LRRK2 inhibition only in postsynaptic AMPAR trafficking. Multiple reports have shown that G2019S does not increase basal Rab10 phosphorylation (Atashrazm et al., 2019; Fan et al., 2018; Liu et al., 2018), thus LRRK2 mutation effects on glutamate release may occur through other substrates. Here, increased pRab10 was observed in VKI mouse brain and cortical cultures, suggesting that VPS35 D620N affects different LRRK2 substrates than G2019S. We did not observe any effect of MLi-2 on VKI presynaptic phenotypes (an effect seen in LRRK2 G2019S scenarios), suggesting they are either LRRK2-independent, or involve LRRK2 GTPase or structural scaffolding domains.

To our knowledge this is the first study of glutamatergic neuron biology in a knock-in model of VPS35 D620N parkinsonism. We found the D620N mutation results in a gain-of-function in GluA1-AMPAR trafficking and glutamate release. LRRK2 kinase inhibition affects post-synaptic AMPAR trafficking in all genotypes, including WT, but has no apparent effect on presynaptic mutant Pr, suggesting caution is necessary in the wide application of LRRK2 kinase inhibitor treatment for PD. We add support to the theory that synaptic transmission is augmented at early time points in PD, which potentially represents early pathophysiological processes that can be targeted to prevent transition to later pathological damage (reviewed in Kuhlmann & Milnerwood, 2020; Picconi, Piccoli, & Calabresi, 2012; Volta, Milnerwood, & Farrer, 2015).

## Materials and methods

### VPS35 D620N knock-in mice and genotyping

Constitutive VPS35 D620N knock-in mice (VKI) were generated by Ozgene (Australia) under guidance of my co-supervisor, Dr. Matthew Farrer, using gene targeting in C57Bl6 embryo stem cells (Bruce4) as previously described (Cataldi et al., 2018). The VKI strain has been deposited in Jackson Labs with open distribution supported by the Michael J Fox Foundation *(VPS35* knock-in: B6(Cg)-Vps35tm1.1Mjff/J). All mice were bred, housed, and handled according to Canadian Council on Animal Care regulations. All procedures were conducted in accordance with ethical approval certificates from the UBC ACC (A16-0088; A15-0105) and the Neuro CNDM (2017-7888B). Animals were group-housed in single-sex cages with littermates after weaning. Single-housed animals were selected as breeding animals or timed pregnancies if possible, and were never used as experimental animals.

Mice were genotyped to allow ongoing colony management, planning, and for *post-mortem* confirmation of the genotype of all experimental animals. The procedure for creating the VKI resulted in 51 base-pair insertion in the non-coding regions of *VPS35*, such that PCR amplification of the WT gene using appropriate primers creates a product of 303 base pairs and the knock-in gene a product of 354 base pairs. The animals were genotyped by PCR amplification of *VPS35*, followed by confirmation of the presence of a 303bp product (WT), a 354bp product (Ho) or both (Het). Small tissue samples (tail tips collected at weaning for colony management, ear tissue collected at euthanasia for *post-mortem* confirmation, or forepaws for primary culture) were digested in 100uL 10% Chelex (Bio-Rad 142-1253) at 95°C for 20 minutes and spun down to result in DNA-containing supernatant. 2uL DNA was mixed with 18uL of master mix containing taq polymerase, buffer (DNAse- and RNAse-free water, 10x buffer, MgCl 25mM, dNTPs 10mM), and primers (ThermoFisher Custom DNA oligos: forward-TGGTAGTCACATTGCCTCTG; reverse-ATGAACCAACCATCAATAGGAACAC) according to the instructions for the taq polymerase kit (Qiagen 201203), and the PCR was performed in a programmable machine (program available upon request). Agarose gel electrophoresis was used to separate the products on a 3-4% gel with fluorescent DNA dye (ZmTech LB-001G) and visualized on a Bio-Rad ultraviolet gel imager.

### Western blots and co-immunoprecipitations

Three-month-old male mice were decapitated, and brains removed and chilled for 1 minute in ice-cold carbogen-bubbled artificial cerebrospinal fluid (ACSF; 125mM NaCl, 2.5mM KCl, 25mM NaHCO3, 1.25mM NaH2PO4, 2mM MgCl2, 2mM CaCl2, 10mM glucose, pH 7.3-7.4, 300-310 mOsm). For region-specific analysis, this was followed by rapid (<6min) microdissection of cortex, striatum, hippocampus, dorsal midbrain, olfactory bulbs, and cerebellum, with all remaining tissue pooled as ‘rest’. Tissues were flash frozen in liquid nitrogen and either lysed for immediate use or stored at −80°C. For *WES* and chemiluminescent western blots, tissues were mechanically homogenized in HEPES buffer (20mM HEPES, 50mM KAc, 200mM Sorbitol, 2mM EDTA, 0.1% Triton X-100, pH 7.2; Sigma Aldrich) containing protease inhibitor cocktail (Roche 11697498001), then incubated on ice for 45 minutes with occasional gentle agitation. For fluorescent western blots, tissue was homogenized by probe sonication at 20kHz for 10s in ice-cold TBS buffer (tris-buffered saline, 1% Triton X-100; pH 7.4) containing protease inhibitor (Roche 11697498001) and phosphatase inhibitor (Sigma 4906845001) cocktails. This buffer was selected to optimize detection of LRRK2 and phosphorylated proteins. Lysate concentrations were quantified by Pierce BCA assay (ThermoFisher 23255) and samples were adjusted to equal concentrations in lysis buffer prior to denaturing.

For traditional chemiluminescence detection, 10-15μg protein was prepared in 4x NuPage LDS sample buffer (Invitrogen NP0008) with 2.5% β-mercaptoethanol or 500mM DTT to a total volume of 10-24uL, and denatured at 70°C for 10 minutes. Samples were loaded into a NuPAge 4-12% Bis-Tris gel (Invitrogen NP0322BOX) in an XCell II Blot module (Invitrogen) and run at 70V for 30 min, followed by 110V for 1h. Separated proteins were transferred to methanol-activated Immobilon-P PVDF membrane (Millipore IPVH00010) for 90 minutes at 25V at room temperature, then blocked with 5% milk in PBS for 1 hour at room temperature. Membranes were probed by shaking with primary antibodies in primary antibody solution (PBS, 2% BSA, 0.05% Tween-20) for either 1 hour at room temperature or overnight at 4°C, washed 4x in PBST (PBS, 0.05% Tween-20), and detected by HRP-conjugated secondary antibodies (Invitrogen; shaken in PBS, 5% milk, 0.05% Tween-20 for 30 minutes at room temperature). Chemiluminescence was detected with Pierce ECL (ThermoFisher 32209) imaged on a Chemi-Doc imaging system (Cell-Bio).

Fluorescence western blots were performed with the following modifications to the above protocol: 1) 40ug protein samples to account for the linear and therefore weaker detection capabilities of fluorescent western blotting; 2) Bolt 4-12% Bis-Tris Plus Gels (Invitrogen NW04120BOX); 3) Fluorescence detection optimized membrane, Immobilon-FL PVDF (Millipore IPFL00010); 4) TBS was substituted for PBS in all buffers to prevent possible cross-reaction of phospho-specific antibodies with phosphate groups; 5) transfer times were increased to 2.5 hours for transfer of larger proteins (LRRK2), and; 6) LiCor fluorescent secondary antibodies were used (LI-COR) and imaged on a LI-COR Odyssey Infrared imaging system (LI-COR).

When necessary, phosphorylated or low-abundance proteins were blotted first, and membranes were stripped using LI-COR NewBlot IR stripping buffer (LI-COR 928-40028; 30m at a time until no signal remained from the first blot), then reblotted for non-phosphorylated or higher abundance proteins of similar sizes.

Images for both types of blots were background-subtracted and analyzed for band intensity with ImageLab software. Signals were normalized to a housekeeping protein quantified from the same gel (GAPDH, β-tubulin, β-actin).

Wherever possible as dictated by equipment availability and antibody compatibility, lysates were blotted on a ProteinSimple *WES* automated capillary-based size sorting system as previously described (Beccano-Kelly, Kuhlmann, et al., 2014). Briefly, lysates were mixed with reducing fluorescent master mix (ProteinSimple SM-001), heated (70°C for 10min) and loaded into manufacturer microplates containing primary antibodies (see below) and blocking reagent, wash buffer, HRP-conjugated secondary antibodies, and chemiluminescent substrate (ProteinSimple DM-001/2). *WES* data was analyzed on manufacturer-provided Compass software.

We used the following primary antibodies: NEEP21/NSG1 (Genscript A01442), FAM21 (Millipore ABT79), VPS35 (Abnova H00055737-M02), VPS26 (a kind gift from J. Bonifacino, NICHD), CI-MPR (a kind gift from M. Seaman, Oxford), GluA1 (Millipore 05-855R), D2R (Millipore AB5084P), GluN1 (Millipore 05-432), LRRK2 (Abcam ab133474), LRRK2 phosphoS935 (Abcam ab133450), Rab10 phosphoT73 (Abcam ab230261), Rab10 (Abcam ab104859), VGluT1 (Millipore AB5905), β-tubulin (Covance MRB-435P), β-Actin (Abcam ab6276), and GAPDH (Cell Signaling 2118; ThermoFisher MA5-15738).

For co-immunoprecipitation, 500μg of protein at 1μg/μL was rotary-incubated overnight at 4°C with VPS35 antibody (Abnova H00055737-M02) or Mouse IgG2a control antibody (Abcam ab18414) coupled to M-280 Tosyl-activated Dynabeads (Invitrogen 14204). Small aliquots of each lysate were set aside before IP to verify the equivalence of starting concentrations. 24h later, loaded beads were washed with ice-cold lysis buffer (x3) prior to resuspension in reducing 1x NuPage LDS sample buffer (for traditional; Invitrogen NP0008) or ProteinSimple fluorescent master mix (for *WES;* ProteinSimple SM-001). Protein was eluted and denatured by heating at 70°C for 10 minutes prior to western blotting as described above.

Antibody specificity was tested using lysate from LRRK2 knock-out mouse brain or Rab10 knock-out AtT20 cell lysates. The LRRK2 knock-out mice have been previously described by Hinkle and colleagues (2012). Rab10 knock-out AtT30 cells were a kind gift from Dr. Peter McPherson, and have been previously described (Alshafie et al., 2020).

### Primary cortical cultures

We mated heterozygous VKI mice to generate embryos used for primary neuronal cultures. Dames were euthanized by rapid decapitation, and embryonic day 16.5 pups were removed and microdissected in Hank’s Balanced Salt Solution (HBSS; Gibco 14170161) with 1x penicillinstreptomycin (penstrep; Sigma-Aldrich P4333) in a petri dish on ice. The cortices from each pup were held in 500uL supplemented Hibernate-E medium (Gibco A1247601; 1x GlutaMax Gibco 35050061; and 1% NeuroCult SM1 StemCell 5711) at 4°C while the pups were genotyped as described above. Genotype-pooled tissue was dissociated chemically for 10min in 0.05% Trypsin-EDTA (Gibco 25300054), followed by deactivation with 10% FBS, then mechanically dissociated in supplemented Neurobasal plating medium (Neurobasal Gibco 21103049; 1x GlutaMax Gibco 35050061; and 1% NeuroCult SM1 StemCell 5711).

Cells were plated onto poly-D-lysine-coated plates or coverslips (Sigma P7280) and matured to 21 days *in-vitro* (DIV21) while incubating at 37°C with 5% CO2. For biochemistry, cells were plated in 2mL medium at 1 million cells/well in 6-well plates. For ICC experiments, cells were plated in 1mL medium onto no.1.5 glass coverslips in 24-well plates. Non-nucleofected cells were plated at 115000 cells/well. For GFP-filled neurons, 1 million cells were nucleofected with 1μg pAAV-CAG-GFP plasmid DNA (Addgene 37825) in Ingenio electroporation buffer (Mirus MIR50111) using an Amaxa Nucleofector2b (Lonza), mixed 1:1 with non-nucleofected cells, and plated at 225k cells/well in 1mL medium as above. From DIV4, 10% fresh media was added to all wells every 3-4 days until use.

### Immunocytochemistry and imaging analysis

Cultured cortical neurons were fixed at DIV21 (4% PFA, 4% sucrose in PBS, 10min), permeabilized where appropriate (ice-cold methanol, 3min), and blocked (5% goat serum in PBS) prior to incubation with primary antibodies for 1 hour at RT or overnight at 4°C. Primary antibodies were prepared in antibody solution (2% goat serum, 0.02% Tween20 in PBS). For surface labelling, the cells were not permeabilized until after primary antibody incubation and no detergent was added to the primary antibodies. Proteins were fluorescently labeled with Alexa-conjugated secondary (Invitrogen) in antibody solution for 30 minutes (RT), and coverslips were slide mounted with Prolong Gold (Invitrogen P36930).

We used the following primary antibodies: GFP (Abcam ab1218); VPS35 (Abnova H00055737); VPS26 (a kind gift from J. Bonifacino, NICHD); FAM21C (Millipore ABT79); NEEP21/NSG1 (Genscript A01442); Rab11 (Abcam ab95375); MAP2 (Abcam ab5392); GluA1 (Millipore 05-855R); PSD95 (Thermo Scientific MA1-045); VGluT1 (Millipore AB5905); GluA1 extracellular (Millipore ABN241); Rab10 (Abcam 237703); and GluA1 (Alomone AGP-009). Rab10 specificity was tested using Rab10 knock-out AtT30 cells (a kind gift from Dr. Peter McPherson; described in (Alshafie et al., 2020)) and cultured cortical neurons (Fig. 8 Supp. 1)

Cells having a pyramidal morphology – triangular or teardrop shaped cell bodies with spiny, clearly identifiable apical and basal dendrites (Kriegstein & Dichter, 1983) – were selected for imaging. All images were blinded and randomized prior to processing and analysis.

Cell density counts were performed manually on DAPI and MAP2 co-labeled images acquired at 10x with an Olympus Fluoview 1000 confocal microscope. Sholl analysis images were acquired at 20x with an Evos FL epifluorescence microscope. Dendritic neurites (excluding axons) were traced and analyzed using the Simple Neurite Tracer plugin for FIJI ImageJ using a radial segmentation of 5 μm.

Images for colocalization were acquired on either an Olympus Fluoview 1000 confocal microscope, with images taken at 60x with 2x optical zoom, in 0.5μm stacks, or a Zeiss Axio Observer with Apotome.2 structured illumination upon which images were taken at 63x in 0.25um stacks. Z-stack acquisition was set to capture all MAP2 stained dendrite for unfilled cells, or GFP-filled dendrite for filled cells. Acquisition parameters were constrained within each culture set. Z-stacks for each channel were flattened using the max projection function in FIJI. MAP2 or GPF stains were used to mask dendrites after their first branch point; primary dendrites and cell bodies were excluded from masks. Areas of the dendritic arbor with many intersecting neurites from other cells were excluded from analyses. Images were manually thresholded to create binary masks of clusters. Briefly, the smallest/dimmest clusters perceived to be above background were reduced to a single pixel, binarized, and then cluster masks re-expanded by a 1 pixel radius in the analysis pipeline. Cluster densities, intensities, areas, colocalization densities, and Pearson’s coefficients were all calculated using automated pipelines in CellProfiler (www.cellprofiler.org; pipelines available upon request). Briefly, the pipeline first uses the dendrite mask to restrict all further analyses to the masked area. From there, the binary masks of clusters are used for the size, density, and colocalization densities within dendrites. Dendrite-masked greyscale images are used for Pearson’s coefficients, and greyscale images are overlaid with the cluster masks to measure intensity within clusters inside the dendritic region selected for analysis.

### Electrophysiology in cell culture

Whole-cell patch-clamp recordings were performed on cortical cells at DIV18-22. Neurons were perfused at room temperature with extra-cellular solution (167mM NaCl, 2.4mM KCl, 1mM MgCl_2_, 10mM glucose, 10mM HEPES, 1mM CaCl_2_, 1μM tetrodotoxin and 100μM picrotoxin, pH 7.4, 290-310mOsm). In MLi-2 experiments, ESC was supplemented with 500nM MLi-2 in 45% Captisol PBS or an equal volume of 45% Captisol PBS.

Pipettes were filled with intracellular solution (130mM Cs-methanesulfonate, 5mM CsCl, 4mM NaCl, 1mM MgCl_2_, 5mM EGTA, 10mM HEPES, 5mM QX-314, 0.5mM GTP, 10mM Na_2_-phosphocreatine, 5mM MgATP, and 0.1mM spermine, at pH 7.3 and 290 mOsm). Pipette resistance was constrained to 3-7MOhms for recording. Recordings were acquired by a Multiclamp 700B amplifier in voltage clamp mode at Vh −70mV, signals were filtered at 2 kHz, digitized at 10kHz. The membrane test function was used to determine intrinsic membrane properties 1 minute after obtaining whole-cell configuration, as described previously (Brigidi et al., 2014; Kaufman et al., 2012; Milnerwood et al., 2012; Munsie et al., 2015).

Tolerance for series resistance was <28 mOhm and uncompensated, and recordings discarded if Rs changed by 10% or more. mEPSC frequency and amplitudes were analyzed with Clampfit 10 (Molecular Devices) with a detection threshold of 5pA and followed by manual confirmation of all accepted peaks; non-unitary events were suppressed from amplitude analysis but retained for frequency.

Peak-scaled nonstationary fluctuation analysis was performed on unitary events from each recording in the following way: events were aligned by peak amplitude, baselines were adjusted, and all events normalized to −1 at their maximum amplitude. Event amplitudes and variance from the mean at each recording interval were calculated using the built-in NSFA plugin in Clampfit 10 (Molecular Devices), then rescaled to pre-normalization values. Mean-variance plots were made in GraphPad Prism using values from the event peak to 5pA above baseline (due to baseline noise), and fit using the least squares method and the second order polynomial function representing the following equation:

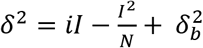

where *δ*^2^ = variance, *i* = single channel current, I = mean current, 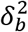 = background variance as previously described by others (T. A. Benke et al., 2001; Hartveit & Veruki, 2007; Smith-Dijak et al., 2019; Traynelis, Silver, & Cull-candy, 1993). For conventional NSFA, *N* = number of open channels at peak current; however the process of peak-normalizing required to analyze mEPSCs renders this value arbitrary. Weighted single-channel conductance was calculated by the following equation:

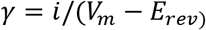

where *γ* = weighted single-channel conductance, *i* = single channel current, *V_m_* = holding potential (−70mV), and *E_rev_* = reversal potential for AMPAR current (0mV in our ECS). Recordings were rejected if the best-fit curve had an R^2^<0.5.

### LRRK2 kinase inhibition with MLi2 treatment

We inhibited LRRK2 kinase activity with the selective LRRK2 inhibitor MLi-2 (Tocris 5756). MLi-2 has low solubility in water, necessitating the use of a vehicle for solubilization. We chose to use the cyclodextrin Captisol (Ligand RC-0C7-100) as a vehicle, due to its worldwide safety approval and use in human drug formulations (www.captisol.com/about). We bath sonicated img MLi-2 in 2mL 45% Captisol-PBS for ~2 hours at room temperature (until complete solubilization of MLi-2). Solutions were filter sterilized prior to use. Primary cortical cultures were treated with 500nM MLi-2 or Captisol-only control (each 1mL well treated with 0.4uL stock in 100mL fresh media; final Captisol concentration 0.00016%) for 2 hours prior to fixation for ICC, lysis for western blot, or transferred to the microscope chamber for electrophysiology experiments. Treatment concentrations and times were selected based on the pharmacokinetic data collected by Fell and colleagues (2015). For western blot experiments in brain tissue, animals were injected intraperitoneally with MLi-2 or Captisol-only control at a dose of 5mg/kg, 2 hours prior to rapid decapitation without anaesthesia. All tissues were collected, frozen, and stored as described above.

### Data visualization and statistics

All statistical analyses and data visualizations were conducted in GraphPad Prism 8. Because biological data is prone to lognormal distribution, outliers were only removed if inspection revealed that they resulted from human error in data collection or processing. Data sets were analyzed for normality using the D’Agostino & Pearson test. In untreated experiments and drug effect comparisons (3 groups) when the data failed the normality test (alpha<0.05), nonparametric tests were used (Kruskal-Wallis with uncorrected Dunn’s post-test). If the data passed the normality test, (alpha>0.05), a parametric test was chosen. When the SDs were not significantly different, a one-way analysis of variance (ANOVA) with uncorrected Fisher’s LSD post-test was used. If the SDs were significantly different, Welch’s ANOVA and an unpaired t-test with Welch’s correction post-test was used. In the event that the n was too low for normality testing, nonparametric tests were used. For Captisol tests (2 groups), the tests chosen were appropriate for two groups (Mann-Whitney for nonparametric and unpaired two-tailed t-test for parametric). MLi-2 treatment data was analyzed using 2-way ANOVA (2-way ANOVA) with uncorrected Fisher’s LSD post-tests.

Sample sizes were selected according to generally accepted standards in the field. No power analyses were conducted to predetermine sample sizes. Data is represented as scatter plots of all data points with mean and standard error of the mean. Sample sizes represent biological replicates. Where n=x(y), x= number of cells imaged/recorded, y=number of independent cultures.

## Supplementary Figures

**Figure 4 supplement 1.**
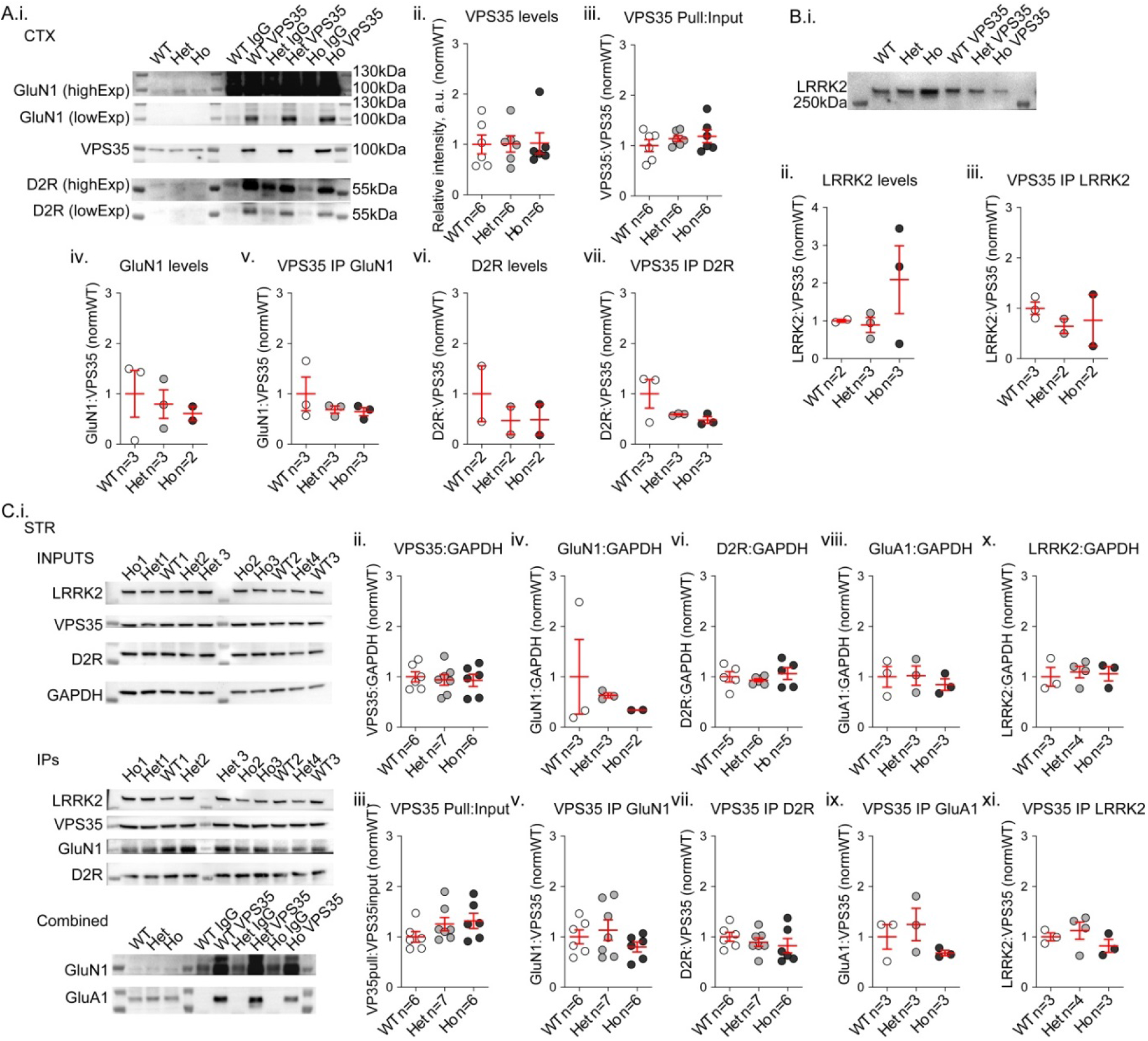
Co-immunoprecipitation in cortical and striatal lysates from 3-month-old mice reveals novel neuronal VPS35 cargoes and no genotype effect on cargo binding. **A)** Western blot of cortical lysates and coIPs were probed for VPS35, D2R, GluA1, GluN1, GluA1, and GAPDH(i). There was no genotype effect on VPS35 levels or IP (ii-iii, 1-way ANOVA *p*>0.99; Kruskal-Wallis *p*=0.62, respectively). NMDA-receptor subunit GluN1 association with retromer has not previously been published; the mutation did not affect GluN1 levels nor coIP with VPS35 (iv-v, Kruskal-Wallis *p*=0.76; *p*=0.44, respectively). D2-type dopamine receptors are a novel cargo; there was no significant genotype effect on D2R levels or CoIP with VPS35 (vi-vii, Kruskal-Wallis *p*=0.80; *p*=0.44, respectively). **B)** CoIP of cortical lysates in A probed for LRRK2 (i). There were no significant genotype effects on LRRK2 levels or association of LRRK2 with VPS35 by coIP (ii-iii Kruskal-Wallis *p*=0.76; *p*=0.52, respectively). **C)** Striatal lysates quantified as in A & B (i). There were no genotype effects on VPS35 levels or pull by the antibody (ii-iii, Kruskal-Wallis *p*=0.97; *p*=0.13, respectively); GluN1 levels or coIP (iv-v, Kruskal-Wallis *p*=0.51; *p*=0.42, respectively); D2R levels or coIP (vi-vii Kruskal-Wallis *p*=0.70; *p*=0.45, respectively); GluA1 levels or coIP (viii-ix, Kruskal-Wallis *p*=0.83; *p*=0.44, respectively); or LRRK2 levels or coIP (x-xi, Kruskal-Wallis *p*>0.99; *p*=0.40, respectively). For all panels, n= number of experimental animals.

**Figure 6 supplement 1.**
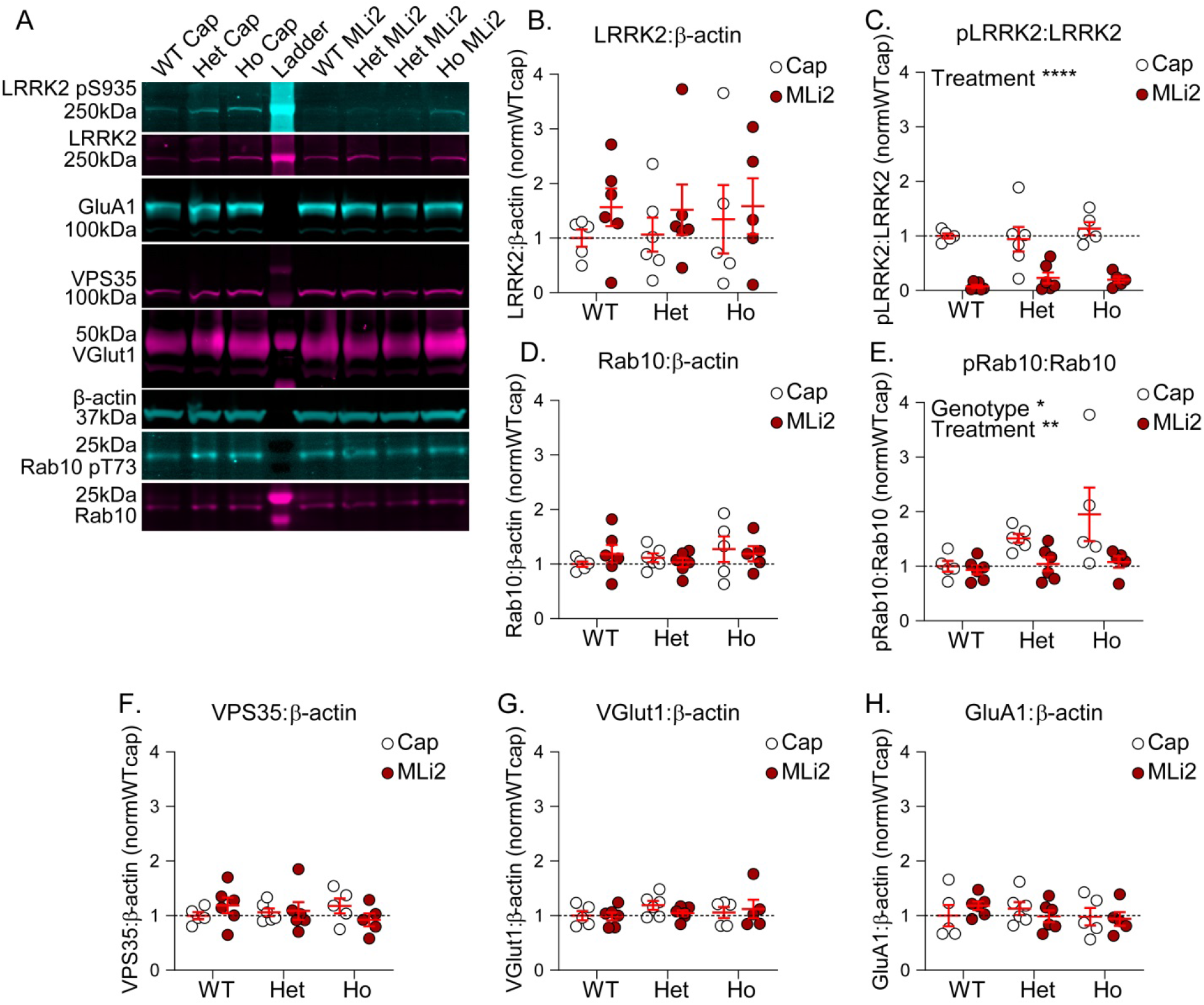
Acute MLi-2 treatment reduces LRRK2 kinase activity in murine brain with no effect on protein expression. **A)** Western blot of whole brain lysate following acute MLi-2 treatment were probed for LRRK2, LRRK2 phospho-S935, GluA1, VPS35, VGluTl, Rab10, Rab10 phospho-T73, and β-actin. **B)** There were no significant effects of genotype or treatment on LRRK2 levels (2-way ANOVA genotype x treatment *p*=0.93; genotype *p*=0.90; treatment *p*=0.24). **C)** ML12 treatment significantly reduced pLRRK2 in all genotypes (2-way ANOVA treatment *p*<0.0001; Uncorrected Fisher’s LSD WT-WTMLi2 \*\*\*\**p*<0.0001; Het-HetMLi2 \*\*\**p*<0.0002; Ho-HoMLi2 \*\*\*\**p*<0.0001). **D-E)** There was similarly no effect of genotype or treatment on Rab10 protein levels (D, 2-way ANOVA genotype x treatment *p*=0.5258; genotype *p*=0.4683; treatment *p*=0.9659), but significant genotype and treatment effects on pRab10 due to significant reductions in homozygous cells (E, 2-way ANOVA genotype *p*=0.04; treatment *p*<0.009; Uncorrected Fisher’s LSD WT-WTMLi2 *p*=0.82; Het-HetMLi2 *p*=0.10; Ho-HoMLi2 \*\**p*=0.007). **F-H)** There were no significant effects of genotype or treatment on protein levels of VPS35 (F, 2-way ANOVA interaction *p*=0.23; genotype *p*=0.94; treatment *p*=0.89), VGluT1 (G, 2-way ANOVA interaction *p*=0.55; genotype *p*=0.39; treatment *p*=0.69), or GluA1 (H, 2-way ANOVA interaction *p*=0.45; genotype *p*=0.61; treatment *p*>0.99). For all experiments, n represents number of experimental animals and are as follows: WTCap n=5, WTMLi2 n=6, HetCap n=6, HetMLi2 n=6, HoCap n=5, HoMLi2 n=5.

**Figure 6 supplement 2.**
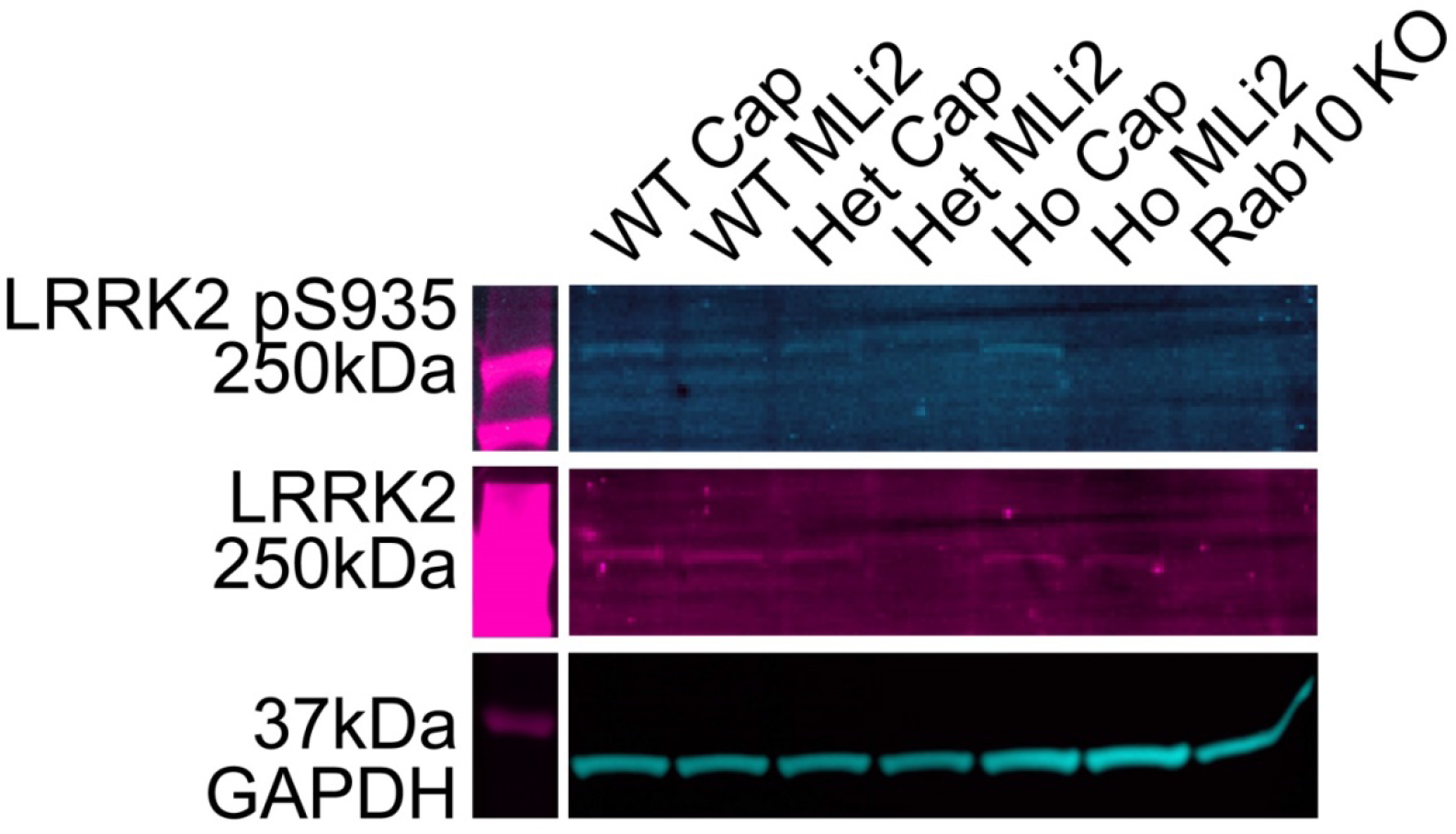
LRRK2 is present and phosphorylated in cultured cortical cells, and shows visible reductions in phosphorylation following acute MLi-2 application. Fluorscence western blot of pLRRK2, LRRK2, and GAPDH VKI cortical culture lysate revealed that the presence of LRRK2 and LRRK2 p935 in vehicle treated cultures, and absence of pLRRK2 following acute MLi-2 treatment; however, due to low stoichiometry bands were not high enough above background to be reliably quantified.

**Figure 8 supplement 1.**
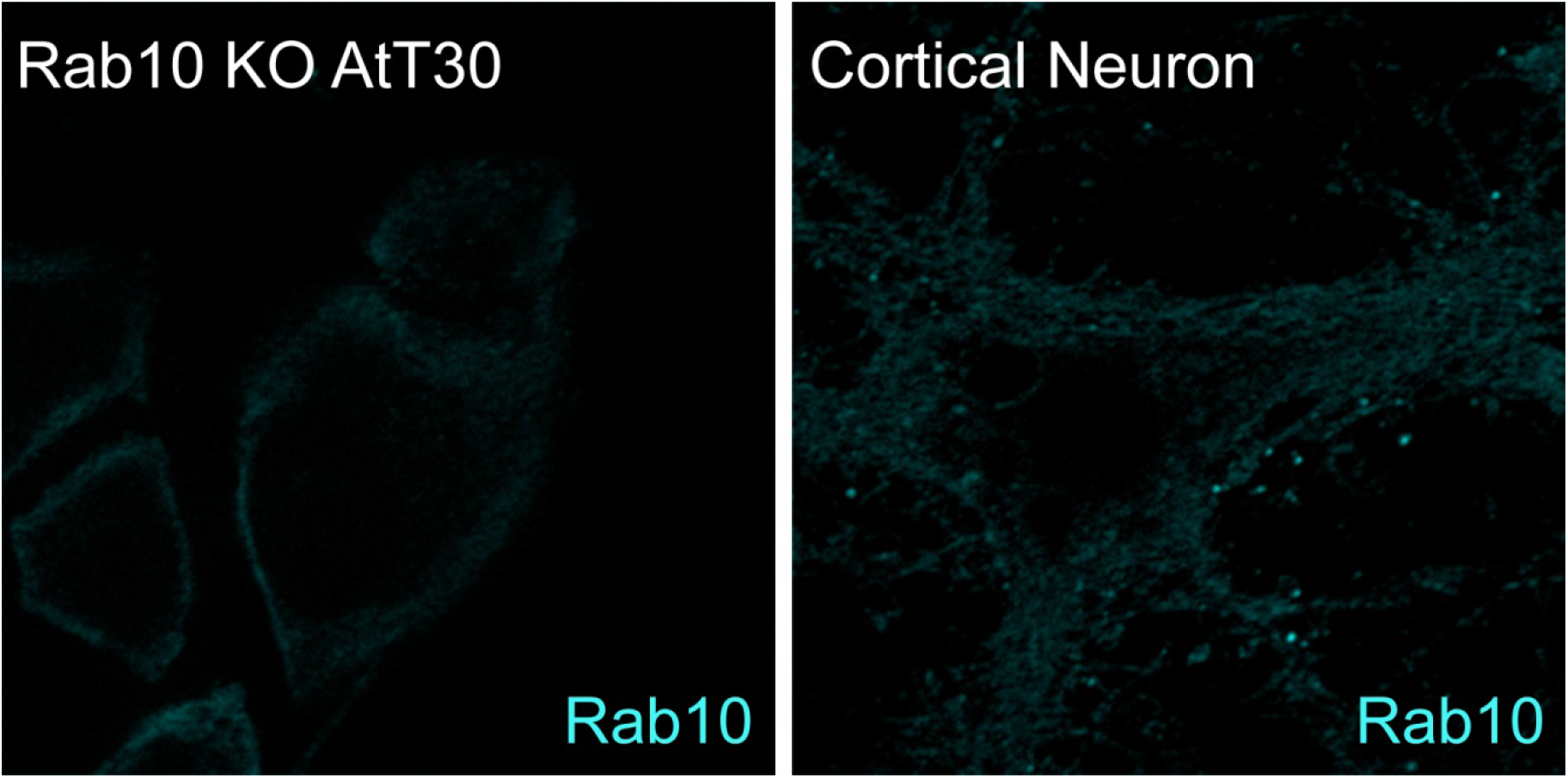
Knock-out testing of specificity of Rab10 antibody for immunocytochemistry. Cultured cortical neurons and Rab10 knock-out AtT30 cells were stained by immunocytochemistry for Rab10, demonstrating punctate staining in the cortical neuron that is absent from the knock-out cells.

